# Mechanism of replication-coupled DNA-protein crosslink proteolysis by SPRTN and the proteasome

**DOI:** 10.1101/381889

**Authors:** Alan Gao, Nicolai B. Larsen, Justin L. Sparks, Irene Gallina, Matthias Mann, Markus Räschle, Johannes C. Walter, Julien P. Duxin

## Abstract

DNA-protein crosslinks (DPCs) are bulky DNA lesions that interfere with DNA metabolism and therefore threaten genomic integrity. Recent studies implicate the metalloprotease SPRTN in S-phase removal of DPCs, but how SPRTN activity is coupled to DNA replication is unknown. Using *Xenopus* egg extracts that recapitulate replication-coupled DPC proteolysis, we show that DPCs can be degraded by SPRTN or the proteasome, which act as independent DPC proteases. Proteasome recruitment requires DPC polyubiquitylation, which is triggered by single-stranded DNA, a byproduct of DNA replication. In contrast, SPRTN-mediated DPC degradation is independent of DPC polyubiquitylation but requires polymerase extension of a nascent strand to the lesion. Thus, SPRTN and proteasome activities are coupled to DNA replication by distinct mechanisms and together promote replication across immovable protein barriers.

**Highlights:**

- The proteasome, in addition to SPRTN, degrades DPCs during DNA replication
- Proteasome-dependent DPC degradation requires DPC ubiquitylation
- DPC ubiquitylation is triggered by ssDNA and does not require the replisome
- SPRTN-dependent DPC degradation is a post-replicative process

## Introduction

DNA-binding proteins perform a variety of functions via transient non-covalent DNA-protein interactions. Exposure to crosslinking agents can cause DNA-binding proteins to become covalently trapped on DNA, forming DNA-protein crosslinks (DPCs). Exogenous crosslinking agents associated with DPC formation include ionizing radiation, UV-light, and chemotherapeutics such as nitrogen mustards, platinum compounds, and topoisomerase and DNA methyltransferase poisons (Barker et al., 2005; Ide et al., 2011; Stingele et al., 2017). Even in the absence of exogenous crosslinking agents, DPCs are commonly occurring lesions caused by the high abundance of aldehydes, reactive oxygen species, and DNA abasic sites in cells (Vaz et al., 2017). While DPCs generated by most crosslinking agents link proteins to uninterrupted duplex DNA (classified as type I DPCs), abortive reactions by topoisomerase I and II form DPCs that are flanked on one side by a single-stranded (type II DPCs) or double-stranded DNA break (type III DPCs), respectively (Barker et al., 2005; Ide et al., 2011; Stingele et al., 2017). Left unrepaired, DPCs stall or inhibit processes such as DNA replication and transcription and thereby threaten genomic integrity (Duxin and Walter, 2015; Ide et al., 2011; Stingele and Jentsch, 2015; Vaz et al., 2017).

Given the frequency and cytotoxicity of DPC lesions, it is not surprising that cells have evolved specific pathways to promote their removal. While several canonical DNA repair pathways such as nucleotide excision repair and homologous recombination have previously been linked to DPC repair (reviewed in (Ide et al., 2011)), recent experiments in yeast identified the metalloprotease Wss1 as a dedicated DPC-repair factor (Stingele et al., 2014b). Wss1 removes DPCs from the genome via direct proteolysis of the cross-linked protein (Balakirev et al., 2015; Stingele et al., 2014b). In contemporaneous experiments, we recapitulated replication-coupled DPC proteolysis in *Xenopus* egg-extracts (Duxin et al., 2014). A type I DPC encountered by the replisome on the leading or lagging strand template is rapidly degraded to a short peptide adduct. Degradation of the DPC facilitates replisome bypass and subsequent DNA synthesis across the lesion site by the translesion synthesis (TLS) polymerase complex Rev1-Polζ (Duxin et al., 2014).

In this manner, the replisome is able to simultaneously overcome DPC lesions and clear them from the genome. Collectively, the experiments in yeast and in *Xenopus* established the existence of a specialized proteolytic DPC-repair pathway, although the protease acting in vertebrates remained elusive at the time.

Studies in mammalian cells suggested that the proteasome participates in DPC removal (Baker et al., 2007; Desai et al., 1997; Lin et al., 2008; Mao et al., 2001; Quiñones et al., 2015; Zecevic et al., 2010). Proteasome inhibition prevents the removal of different types of DPCs including trapped topoisomerases I and II and DNA polymerase β (Desai et al., 1997; Lin et al., 2008; Mao et al., 2001; Quiñones et al., 2015), and sensitizes cells to formaldehyde treatment (Ortega-Atienza et al., 2015). Additionally, DPC polyubiquitylation was observed in the case of covalent topoisomerase I cleavage complexes (Desai et al., 1997). However, polyubiquitylation of the more abundant type I DPCs has not been reported (Nakano et al., 2009), and it is therefore uncertain whether DPCs are generally targeted by the proteasome (Vaz et al., 2017). In *Xenopus* egg extracts, inhibition of the proteasome on its own does not significantly stabilize type I DPCs during DNA replication (Duxin et al., 2014). Therefore, whether the proteasome acts on different types of DPCs and whether this process operates during DNA replication remain open questions.

Recently, the metalloprotease SPRTN (also known as Spartan or DVC1) has been implicated in DPC degradation in higher eukaryotes (Lopez-Mosqueda et al., 2016; Maskey et al., 2017; Morocz et al., 2017; Stingele et al., 2016; Vaz et al., 2016). SPRTN shares homology with the yeast DPC protease Wss1 and is proposed to be functionally similar, although whether these two proteins are evolutionarily related remains controversial (Stingele et al., 2014a; Vaz et al., 2017). In humans, mutations in SPRTN that compromise SPRTN’s protease activity cause Ruijs-Aalfs Syndrome (RJALS), which is characterized by genomic instability, premature aging, and early onset hepatocellular carcinoma (Lessel et al., 2014). In mice, loss of SPRTN is embryonically lethal, and conditional inactivation of SPRTN in MEFs blocks cell proliferation (Maskey et al., 2014). Although SPRTN was initially characterized as a regulator of translesion DNA synthesis (Centore et al., 2012; Davis et al., 2012; Mosbech et al., 2012), several reports now suggest that its essential role in cellular viability and genome maintenance involves DPC proteolysis (Lopez-Mosqueda et al., 2016; Maskey et al., 2017; Morocz et al., 2017; Stingele et al., 2016; Vaz et al., 2016). SPRTN is predominantly expressed in S phase and associates with various replisome components (Ghosal et al., 2012; Kim et al., 2013; Mosbech et al., 2012; Vaz et al., 2016). In the absence of SPRTN protease activity, cells accumulate DPCs and exhibit severely impaired replication fork progression (Lessel et al., 2014; Morocz et al., 2017; Vaz et al., 2016). Collectively, these data indicate that DPCs readily form *in vivo* and that cells depend on SPRTN-dependent DPC removal to suppress genome instability, cancer, and aging.

A protease that degrades DPCs during DNA replication must be carefully regulated to avoid nonspecific degradation of replisome components, which would have catastrophic consequences for cells. Indeed, SPRTN proteolytic activity appears to be regulated via different mechanisms. First, SPRTN undergoes monoubiquitylation (Mosbech et al., 2012), which is proposed to prevent its recruitment to chromatin (Stingele et al., 2016). DPC induction triggers SPRTN deubiquitylation by an unknown ubiquitin protease, allowing SPRTN to localize to chromatin and initiate DPC degradation (Stingele et al., 2016). Once SPRTN is recruited to chromatin, DNA binding stimulates its protease activity (Lopez-Mosqueda et al., 2016; Morocz et al., 2017; Stingele et al., 2016; Vaz et al., 2016), and some evidence indicates that SPRTN is uniquely activated by ssDNA (Stingele et al., 2016). SPRTN also degrades itself, which might help switch off its proteolytic function when repair is complete (Stingele et al., 2017). In conjunction, these different layers of regulation offer some explanation for how SPRTN activity is held in check. However, they do not explain how SPRTN activity is directed to DPCs during DNA replication.

Here, we investigated the molecular mechanisms that link DPC degradation to DNA replication. We report that SPRTN and the proteasome function as two independent DPC proteases that operate during DNA replication. Proteasome recruitment to DPCs is strictly dependent on DPC polyubiquitylation, which is triggered by the ssDNA generated during DNA replication. In contrast SPRTN-dependent DPC degradation does not require DPC ubiquitylation, but instead is triggered by the extension of a nascent strand to the DPC. Our results unravel how SPRTN and proteasome activities are targeted to DPCs to facilitate replication across these covalent barriers.

## Results

### DPCs are ubiquitylated and degraded during DNA replication

To investigate DPC repair, the DNA methyltransferase HpaII (M.HpaII, 45 kDa) was trapped at a fluorinated recognition site on a plasmid (Chen et al., 1991). During replication of the resulting plasmid (pDPC) in *Xenopus* egg extracts, converging forks transiently stall at the DPC, after which daughter plasmid molecules are resolved (Figure S1A; (Duxin et al., 2014)). While the daughter molecule containing the undamaged parental strand immediately accumulates as a supercoiled plasmid, the daughter molecule containing the DPC initially migrates as an open circular (OC) species, and is then gradually converted to a supercoiled (SC) repair product through proteolysis of the DPC and translesion DNA synthesis across the resulting peptide adduct (Figures S1A and S1B; (Duxin et al., 2014)). To monitor the integrity of the DPC during replication, we pulled down the plasmid under stringent conditions that disrupt non-covalent nucleoprotein complexes, digested the DNA, and analyzed M.HpaII via Western blotting (Figure 1A). At the 15 minute time point, when replication was well underway (Figure S1B), the covalently attached M.HpaII migrated as a ladder of slow mobility species that subsequently disappeared (Figure 1B, lanes 2-4). Addition of FLAG-ubiquitin to the extract shifted the mobility of the M.HpaII species (Figure 1C, compare lanes 2-3 and 4), indicating that they correspond to ubiquitylated M.HpaII. This conclusion was confirmed by immunoprecipitating the same reaction with anti-FLAG antibody and blotting against M.HpaII (Figure 1D). When DNA replication initiation was blocked with Geminin (Tada et al., 2001; Wohlschlegel et al., 2000), M.HpaII persisted in a largely unmodified form (Figure 1B, lanes 5-6), demonstrating that DPC ubiquitylation and degradation are dependent on DNA replication. In the absence of DNA replication, different modified M.HpaII species slowly appeared (Figure 1B, lane 6). These species were cleaved by the SUMO protease Ulp1 but not the ubiquitin protease USP2 (data not shown), and their appearance was dependent on the SUMO ligase UBC9 (Figure S1C). In contrast, the replication-dependent species did not involve SUMOylation (Figure S1D). Therefore, DPCs undergo both replication-dependent ubiquitylation, which contributes to proteolysis (see below), and replication-independent SUMOylation, whose function remains to be determined.

**Figure 1.**
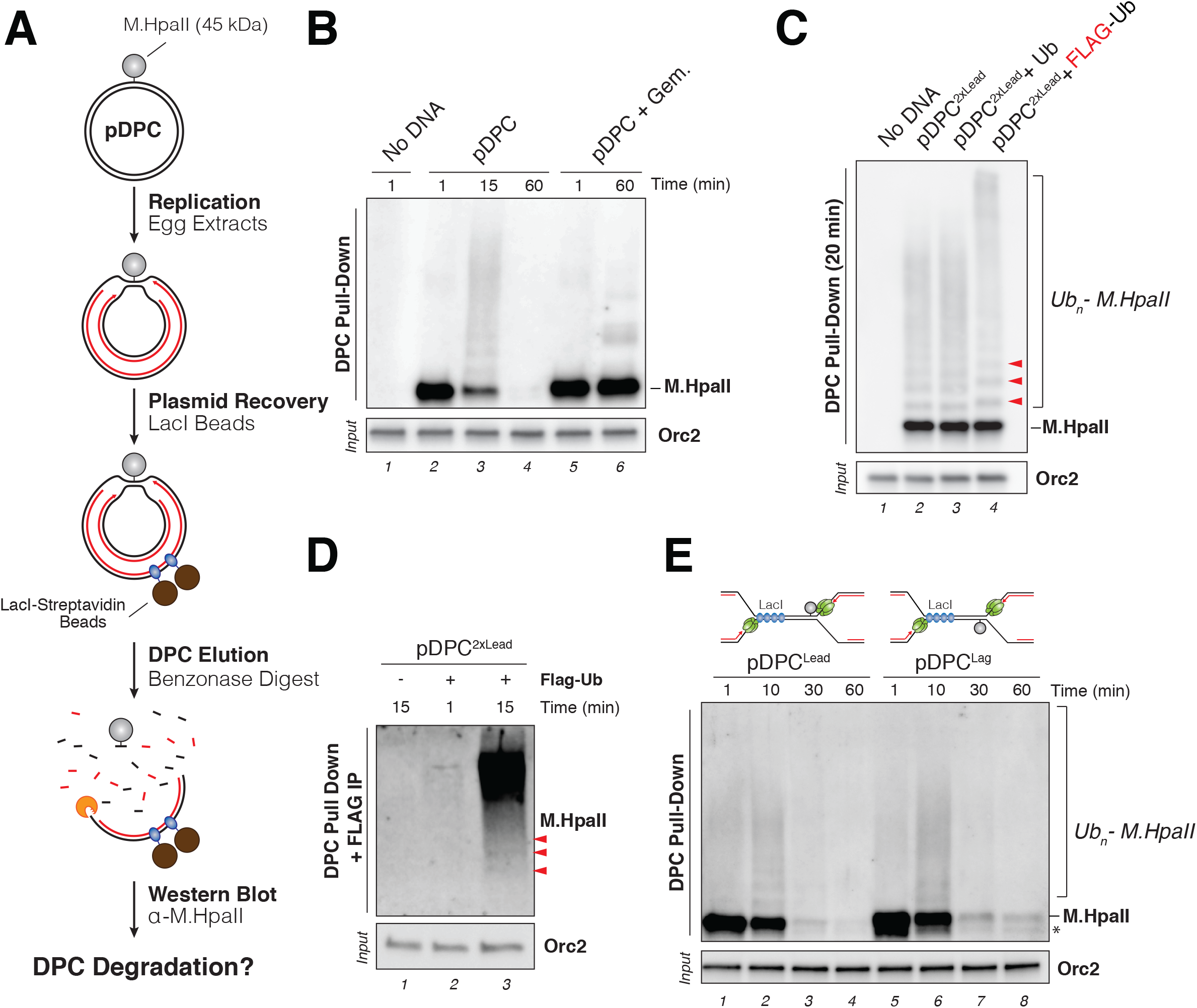
Replication-coupled ubiquitylation and degradation of a DNA-protein crosslink. (**A**) Schematic of the DPC recovery assay. (**B**) pDPC was replicated in egg extracts. Geminin (+ Gem.) was added where indicated to block DNA replication. DPCs were recovered as illustrated in (A) at the indicated time points, and DPCs were blotted with a M.HpaII antibody. Input samples were blotted with a ORC2 antibody. (**C**) pDPC^2xLead^, a plasmid containing two DPCs (one on each strand, see Figure 2A) was replicated in egg extracts supplemented with free ubiquitin (Ub) or FLAG-ubiquitin where indicated. At 20 minutes, DPCs were recovered and blotted with M.HpaII antibody as in (B). Red arrowheads indicate the mobility shift induced by FLAG-ubiquitin on mono-, di-, and tri-ubiquitylated M.HpaII. (**D**) pDPC^2xLead^ was replicated in egg extract supplemented with FLAG-ubiquitin where indicated. DPCs were recovered as in (B) and subsequently immunoprecepitated with anti-FLAG-resin. Ubiquitylated DPCs were detected with M.HpaII antibody. Red arrowheads indicate location of mono-, di-, and tri-ubiquitylated M.HpaII. (**E**) pDPC^Lead^ or pDPC^Lag^ were replicated in egg extract in the presence of LacI to ensure that a single replication fork encounters the DPC (depicted in the upper schemes; (Duxin et al., 2014)). Recovered DPCs were blotted against M.HpaII as in (B). * indicates residual uncrosslinked M.HpaII.

Replication forks promote destruction of DPCs encountered on both the leading and lagging strand templates (Duxin et al., 2014). However, because the replicative CMG helicase translocates on the leading strand template (Fu et al., 2011), it is possible that DPCs encountered on the leading and lagging strands undergo different processing. To address this, we replicated a plasmid containing a lac repressor array that is flanked on one side by a DPC on the top or bottom strand (Figure 1E). The rightward fork stalls at the array in both templates, whereas the leftward fork encounters the DPC on the leading or lagging strand template, respectively (Dewar et al., 2015; Duxin et al., 2014). As shown in Figure 1E, DPCs encountered on either strand were ubiquitylated and degraded with similar kinetics, suggesting that leading and lagging strand DPCs are recognized and processed similarly.

Previously, we demonstrated that DPC degradation is drastically inhibited by ubiquitin-vinyl-sulfone (UbVS), which inhibits most DUBs and therefore depletes free ubiquitin in egg extracts (Dimova et al., 2012; Duxin et al., 2014). We confirmed this result by directly monitoring M.HpaII in UbVS-treated extracts via DPC pull-down. As shown in Figure S1E, UbVS noticeably delayed ubiquitylation of M.HpaII and strongly inhibited DPC proteolysis (lanes 7-11), effects that were rescued with free ubiquitin (lanes 12-16). Collectively, these experiments demonstrate that when a replication fork encounters a DPC on the leading or the lagging strand template, the DPC undergoes extensive polyubiquitylation prior to being degraded.

### SPRTN and the proteasome accumulate on replicating DPC plasmids

To identify DPC protease(s), we combined plasmid pull-down with quantitative high-resolution mass spectrometry (Figure 2A). As shown below, this new approach, which we call plasmid pull down mass spectrometry (PP-MS), faithfully reports the time-resolved accumulation of proteins on plasmids as they undergo replication and repair in *Xenopus* egg extracts. In contrast to CHROMASS, which detects proteins on randomly damaged sperm chromatin (Räschle et al., 2015), PP-MS facilitates identification of proteins associated with defined DNA lesions and discrete repair intermediates (Figure 2A). To validate PP-MS, we first incubated an undamaged control plasmid (pCTRL) in egg extracts and isolated it at the peak of DNA replication at 10 minutes or after replication was completed at 40 minutes (Figure 2A, autoradiograph, lanes 1-3). As expected, CMG, all three replicative DNA polymerases, and most known components of the replication progression complex, were significantly enriched at 10 minutes when replication was ongoing (Figures 2B, columns 1-2, green factors, and S2A). A similar enrichment of replisome components was observed when the plasmid recovered at 10 minutes was compared to a reaction supplemented with Geminin, which inhibits replication initiation (Figures 2B, compare columns 1 and 3, S2A, and S2B). We conclude that PP-MS is a robust method to detect proteins associated with plasmids in egg extracts.

**Figure 2.**
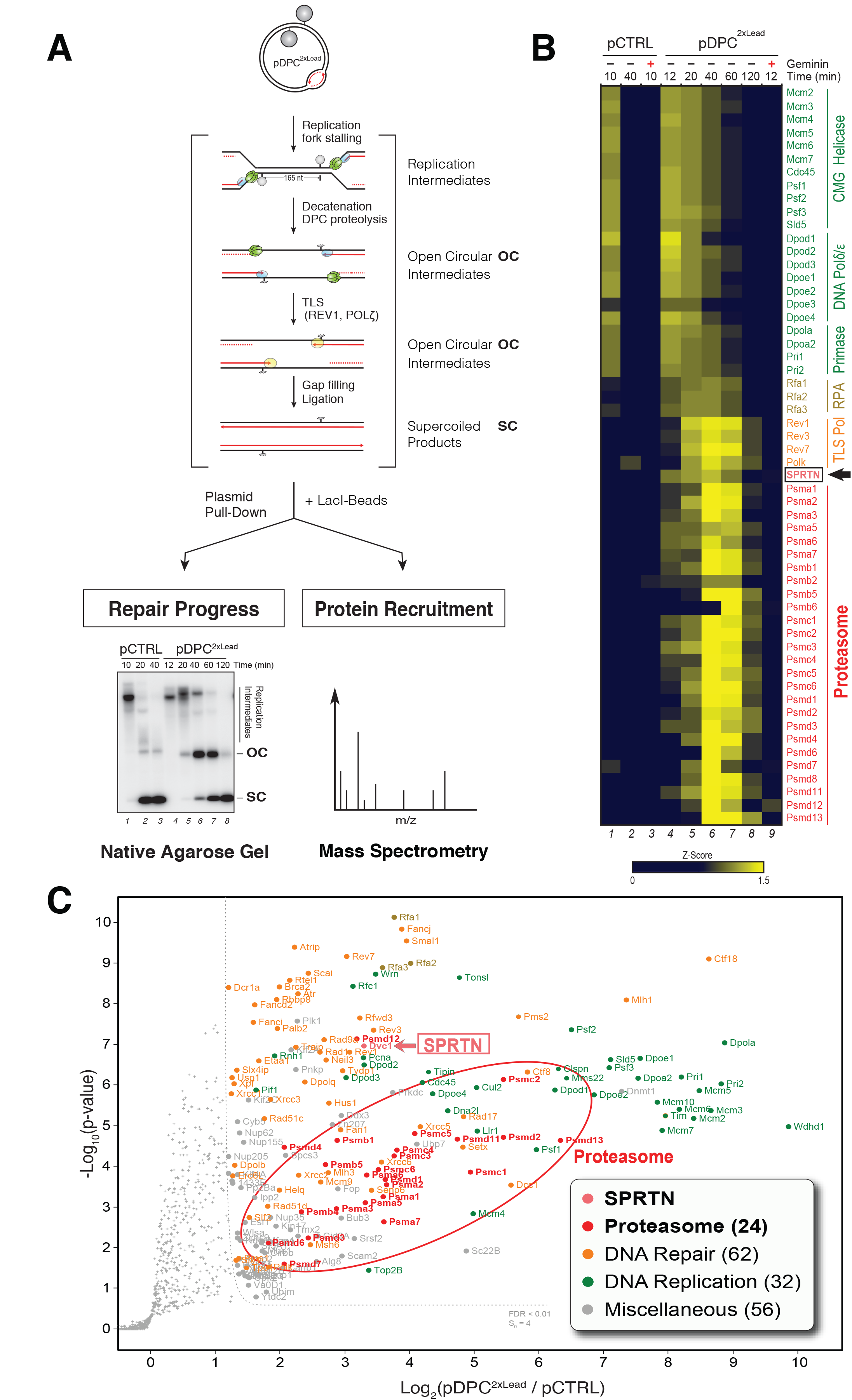
SPRTN and the proteasome are recruited to a DPC plasmid during replication. (**A**) Depiction of replication, recovery, and analysis of pDPC^2xLead^. The cartoon shows how two gapped open circular (OC) daughter molecules are generated upon decatenation of the replication intermediates. Upon degradation of the DPC, translesion synthesis and gap filling restores two supercoiled circular (SC) products. To monitor the progress of the repair reaction, pDPC^2xLead^ was replicated in the presence of [α-^32^P]dATP and replication intermediates were analyzed by agarose gel electrophoresis. In parallel, plasmids were isolated together with the bound proteins by a LacI pull-down as in (Budzowska et al., 2015) and analyzed by label-free mass spectrometry. (**B**) Heat map showing the mean of the z-scored log_2_ LFQ intensity from 4 biochemical replicates. In green, core replisome components; in brown, RPA; in orange, TLS factors; in red, SPRTN and 26S proteasome subunits. (**C**) Analysis of protein recruitment to pDPC^2xLead^ compared to pCTRL. Both plasmids were recovered at 40 min. The volcano plot shows the mean difference of the protein intensity plotted against the p-value calculated by a modified, one-sided T-Test. Full results are reported in Tables S1 and S2.

We next applied PP-MS to DPC repair. To maximize the yield of DPC repair factors, we replicated pDPC^2xLead^, a plasmid containing two DPCs positioned 165 nt apart such that both converging forks encountered a DPC on the leading strand template (Figure 2A). Following fork stalling at the DPC, the daughter molecules underwent decatenation, and the open circular plasmids were repaired by translesion DNA synthesis (Figures 2A, cartoon depiction and autoradiograph, lanes 4-8). Consistent with replication fork stalling at leading strand DPCs (Duxin et al., 2014; Fu et al., 2011), replisome components persisted for up to 40 minutes on pDPC^2xLead^ (Figures 2B, columns 4-8, and S2C, MCM6 panel). The TLS polymerases REV1, Pol ζ and Polκ were recruited to pDPC^2xLead^ following replisome unloading (Figure 2B, orange factors, and S2C, REV1 panel), and their peak binding correlated with the transition from OC to SC plasmid (Figure 2A, lanes 6-7), which depends on the REV1-Polζ complex (Duxin et al., 2014). By 120 minutes, when all molecules had undergone replication-coupled DPC repair (Figure 2A, lane 8), repair factors were largely lost from DNA (Figures 2B, column 8). Besides these replisome and TLS factors, numerous proteins involved in replication-coupled repair processes such as homologous recombination also specifically accumulated on replicating DPC plasmids (see Tables S1 and S2).

Consistent with the recent demonstration that SPRTN functions in mammalian S phase DPC repair (Lopez-Mosqueda et al., 2016; Maskey et al., 2017; Morocz et al., 2017; Stingele et al., 2016; Vaz et al., 2016), we observed a specific enrichment of SPRTN on replicating pDPC^2xLead^ (Figure 2B, columns 4-8). SPRTN recruitment occurred during the peak of proteolysis (20-60 minutes), depended on DNA replication, and was not detected on pCTRL (Figures 2B, 2C, S2C, and S2D). Strikingly, the 26S proteasome was also specifically enriched on DPC plasmids. Out of 33 subunits that form the 26S proteasome, 26 showed significant enrichment on pDPC^2xLead^ compared to pCTRL (Figure 2B, red factors). Proteasome recruitment also peaked between 20 and 60 minutes, was specific to the DPC, and depended on DNA replication (Figures 2B, 2C, S2C, and S2D). Collectively, these experiments provide an unbiased resource of candidate DPC repair factors and single out SPRTN and the proteasome as two proteases that might mediate DPC destruction in egg extracts. They also illustrate the ability of PP-MS to identify proteins associated with different stages in the repair of a chemically-defined DNA lesion.

### SPRTN participates in replication-coupled DPC proteolysis

We first investigated the role of SPRTN in cell-free DPC repair. We immunodepleted SPRTN from egg extracts (Figures 3A and S3A) and examined the effect on pDPC replication. Compared to mock-depleted extracts, which supported rapid accumulation of supercoiled products, SPRTN-depleted extracts exhibited a slight persistence of open circular intermediates (Figures 3B, compare lanes 3-4 with 8-9, red arrowheads, and 3F, lanes 6-10). This defect was reversed by recombinant wild type (WT) SPRTN but not catalytically inactive (EQ) SPRTN (Figure 3B, lanes 11-15 and 16-20) (Stingele et al., 2016). Addition of SPRTN-EQ not only failed to rescue SPRTN depletion but caused a further stabilization of open circular repair intermediates. This inhibitory effect was also observed after addition of excess SPRTN-EQ to undepleted extracts (Figures S3B and S3C, lanes 6-10). Our results indicate that SPRTN deficiency causes a small delay in cell-free DPC repair, and that inactive SPRTN dominantly inhibits repair.

**Figure 3.**
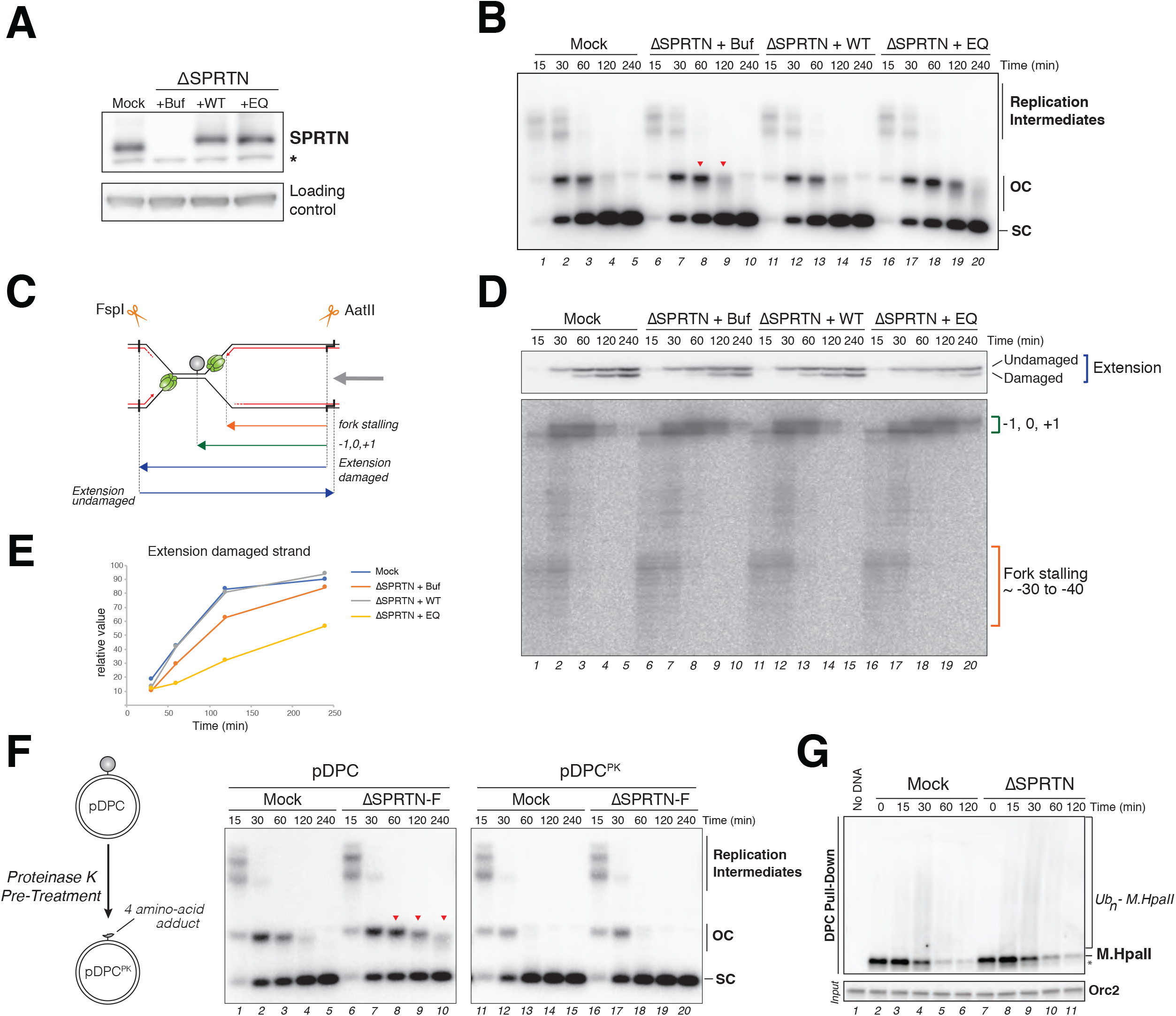
SPRTN protease participates in replication-coupled DPC repair. (**A**) Mock-depleted and SPRTN-depleted egg extracts were blotted with SPRTN and MCM6 (loading control) antibodies. SPRTN-depleted extracts were supplemented with either buffer (+buf), recombinant FLAG-SPRTN (+WT), or recombinant catalytically inactive FLAG-SPRTN E89Q (+EQ). (**B**) The extracts from (A) were used to replicate pDPC in the presence of [α-^32^P]dATP. Samples were analyzed by native agarose gel electrophoresis. Note that open circular molecules that accumulate in egg extracts are subjected to 5’ to 3’ end resection and smear down on the gel (lanes 9, 19-20 and (Duxin et al., 2014)). OC, open circular; SC, supercoiled. Red arrowheads indicate the accumulation of OC molecules. (**C**) Depiction of nascent strands generated after AatII digestion of pDPC. Extension products are monitored with FspI and AatII digestion. CMG helicase is depicted in green and nascent strands in red. Note that digestion with FspI and AatII yields extension products of the damaged and undamaged strand that differ 4 nt in size. (**D**) Samples from (B) were digested with FspI and AatII and separated on a denaturing polyacrylamide gel. Nascent strands generated by the leftward fork are indicated in brackets (lower panel). Extension products of the undamaged and DPC-containing strands are shown in the upper panel. 0 position denotes location of the crosslink. Note that the “-1, 0, +1” stalling positions were erroneously annotated as “0, +1, +2” in (Duxin et al., 2014); see Figure S3G. (**E**) Quantification of the extension of the damaged strand from (D). Values were normalized to the extension of the undamaged strand. (**F**) Left scheme depicts the generation of pDPC^PK^ via Proteinase K treatment of pDPC. pDPC and pDPC^PK^ were replicated in mock-depleted or SPRTN-depleted extracts. Samples were analyzed as in (B). ΔSPRTN-F denotes depletion with an antibody raised against a protein fragment of SPRTN (Figure S3A and material and methods). Red arrowheads indicate the accumulation of OC molecules. (**G**) pDPC was replicated in mock-depleted and SPRTN-depleted extracts. DPCs were recovered and monitored by blotting against M.HpaII as in Figure 1B.

To determine how SPRTN depletion affects replication across a DPC lesion, we digested pDPC replication intermediates with FspI and AatII (Figure 3C) and analyzed nascent leading strands of the leftward fork on a denaturing polyacrylamide gel (Figure 3D) (Duxin et al., 2014). In mock-depleted extracts, the nascent leading strand stalled ~30-40 nucleotides from the DPC due to CMG helicase stalling at the DPC (“fork stalling”) and again directly at the lesion site (“-1, 0, +1”), before progressing past the lesion (“extension”) (Figure 3D, lanes 1-5). SPRTN depletion had no effect on the approach of nascent leading strands to the DPC, but resulted in prolonged stalling at the −1, 0, +1 positions and a corresponding delay in extension on the damaged template strand (Figures 3D, lanes 6-10, and 3E). This effect was rescued by SPRTN-WT but not SPRTN-EQ (Figures 3D, lanes 11-15 and 16-20, and 3E). Together, these experiments demonstrate that SPRTN protease activity facilitates DNA replication across a DPC lesion. Importantly, the inhibitory effect of SPRTN depletion was neutralized by pre-treatment of pDPC with Proteinase K, which is predicted to reduce M.HpaII to a four amino acid peptide adduct (Figures 3F, lanes 16-20 and S3D). Proteinase K pre-treatment also eliminated the dominant negative effect caused by SPRTN-EQ (Figure S3C). However, pDPC^PK^ replication was still blocked when TLS was inhibited via REV1 depletion (Figures S3E and S3F). We conclude that the main function of SPRTN in replication-coupled DPC repair involves DPC proteolysis, which facilitates TLS past the adduct. Nevertheless, SPRTN depletion only modestly preserved M.HpaII levels in the DPC recovery assay (Figure 3G). This result suggests that in the absence of SPRTN, DPCs are still degraded, likely by a second protease.

### Both SPRTN and the proteasome degrade DPCs during replication

Given its recruitment to replicating DPC plasmids (Figure 2B), the proteasome might promote DPC proteolysis in the absence of SPRTN. To test this possibility, we first replicated pDPC^2xLead^ in the presence of the proteasome inhibitor MG262 and observed no delay in conversion of open circular intermediates to supercoiled repair products (Figure 4A, lanes 6-10), as previously seen for pDPC (Duxin et al., 2014). However, in combination with SPRTN depletion, MG262 increased the persistence of gapped repair intermediates (Figure 4A, lanes 16-20), indicating that in the absence of both proteasome and SPRTN activities, DPC repair is severely inhibited. Accordingly, whereas either SPRTN depletion or MG262 treatment alone resulted in only a modest delay in DPC degradation (Figure 4B, lanes 6-8 and 9-11), combined proteasome inhibition and SPRTN depletion greatly stabilized ubiquitylated M.HpaII species (Figure 4B, lanes 12-14). Importantly, DPC degradation was restored by the addition of recombinant SPRTN-WT but not SPRTN-EQ (Figure 4C). Finally, we confirmed the role of the proteasome via immunodepletion with antibodies against the PSMA1 proteasome subunit (Figure S4A), which closely resembled MG262 treatment (Figures S4B-C).

**Figure 4.**
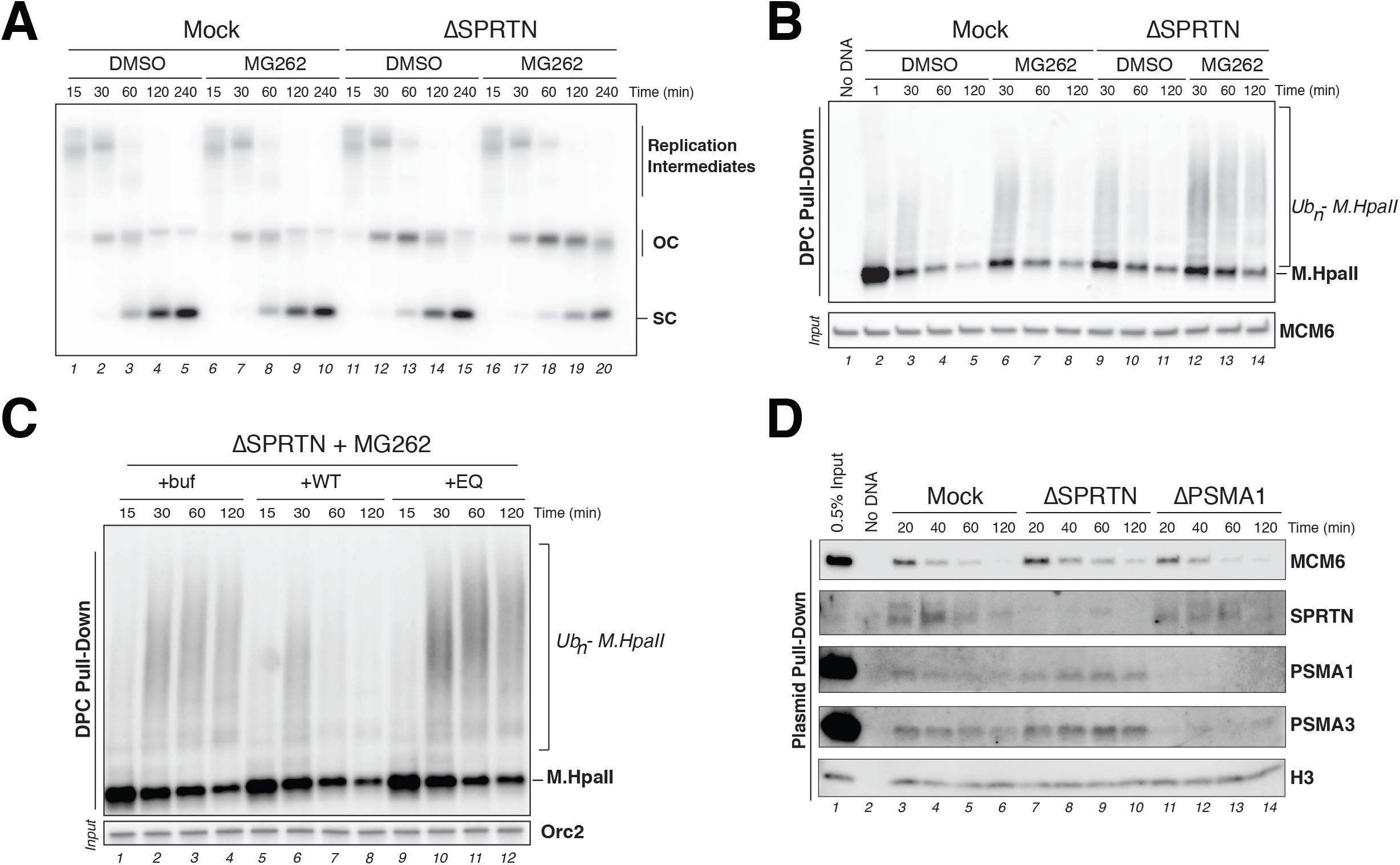
SPRTN and the proteasome degrade polyubiquitylated DPCs. (**A**) Mock-depleted and SPRTN-depleted extracts were used to replicate pDPC^2xLead^. 200 μM of MG262 was added where indicated. Samples were analyzed as in Figure 3B. (**B**) DPCs from (A) were recovered and monitored by blotting against M.HpaII as in Figure 1B. (**C**) SPRTN-depleted egg extracts supplemented with 200 μM of MG262 were used to replicate pDPC^2xLead^. Extracts were supplemented with either buffer (+buf), SPRTN WT (+WT) or SPRTN protease dead (+EQ). DPCs were monitored as in Figure 1B. (**D**) Mock-depleted, SPRTN-depleted or PSMA1-depleted extracts were used to replicate pDPC^2xLead^. Plasmids were recovered as depicted in in Figure 2A and protein-recruitment to the plasmid was monitored with the indicated antibodies (Budzowska et al., 2015).

Given that depletion of SPRTN or inhibition of the proteasome did not prevent DPC degradation, we hypothesized that these two proteases are recruited to a DPC lesion independently of one another. To test this idea, we used plasmid pull-down to monitor the chromatin recruitment of these proteases during replication of pDPC^2xLead^ in extracts depleted of either SPRTN or the proteasome. As shown in Figure 4D, neither SPRTN nor proteasome depletion impaired the recruitment of the other protease to chromatin. In fact, proteasome recruitment was modestly increased in the absence of SPRTN (Figure 4D, lanes 7-10). We conclude that during DNA replication, both SPRTN and the proteasome can degrade DPCs independently of each other.

### SPRTN, but not the proteasome, can degrade non-ubiquitylated DPCs

To investigate the role of DPC ubiquitylation in DPC proteolysis, we chemically methylated the lysines of M.HpaII before conjugating it to the plasmid (Walter et al., 2006), thereby generating a DPC that cannot be ubiquitylated (me-DPC). As shown in Figure 5A, methylated M.HpaII recovered from replication reactions migrated as a single, unmodified band, reflecting a block of DPC ubiquitylation (lanes 8-13). However, even in the absence of ubiquitylation, M.HpaII levels on the plasmid slowly decreased, and a M.HpaII degradation product of ~34 kDa accumulated (Figure 5A, lanes 10-13). In plasmid pull-downs, M.HpaII methylation abolished proteasome recruitment while SPRTN recruitment was only slightly attenuated (Figure 5B, lanes 9-14), suggesting that SPRTN but not the proteasome can still act on the methylated DPC. Consistent with this idea, SPRTN depletion completely stabilized the methylated M.HpaII and abolished formation of the M.HpaII degradation fragment (Figure 5C, lanes 6-10). This defect was reversed by SPRTN-WT but not catalytically inactive SPRTN-EQ (Figure 5D, lanes 6-7 and 14-15). Conversely, MG262 had no effect on me-DPC proteolysis or formation of the M.HpaII fragment (Figures S5A-C). We conclude that SPRTN but not the proteasome can degrade non-ubiquitylated DPCs.

**Figure 5.**
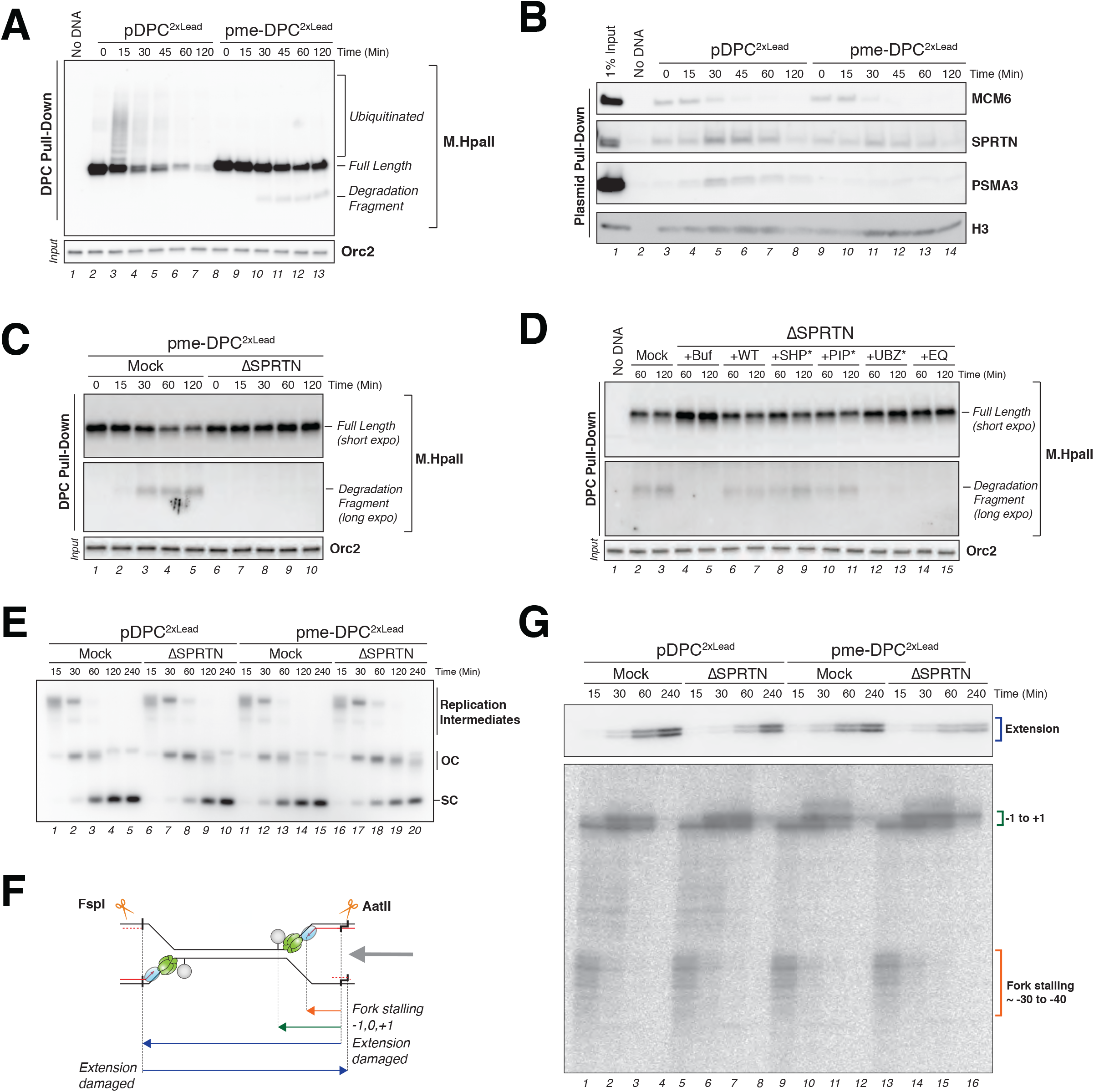
SPRTN, but not the proteasome, can degrade non-ubiquitylated DPCs. (**A**) pDPC^2xLead^ and pme-DPC^2xLead^ were replicated in egg extracts and DPCs monitored like in Figure 1B. Note the concomitant disappearance of full length M.HpaII and appearance of degradation product during replication of pme-DPC^2xLead^. (**B**) pDPC^2xLead^ and pme-DPC^2xLead^ were replicated in egg extracts and indicated protein recruitment to the plasmid was monitored like in Figure 4D. (**C**) pme-DPC^2xLead^ was replicated in mock-depleted or SPRTN-depleted egg extracts. DPCs were recovered and monitored like in Figure 1B. Both a long and a short exposure of the M.HpaII blot are shown where indicated. (**D**) pme-DPC^2xLead^ was replicated in mock-depleted and SPRTN-depleted extracts. SPRTN-depleted extracts were supplemented with either buffer (+buf), or recombinant FLAG-SPRTN variants (for details about the corresponding mutations see Figure S5E). DPCs were recovered and monitored like in Figure 1B. Both a long and a short exposure of the M.HpaII blot are shown where indicated. (**E**) pDPC^2xLead^ and pme-DPC^2xLead^ were replicated in mock-depleted or SPRTN-depleted extracts. Samples were analyzed as in Figure 3B. (**F**) Schematic depicting nascent leading strands and extension products generated by FspI and AatII digest during replication of pDPC^2xLead^ and pme-DPC^2xLead^. (**G**) Samples from (E) were digested with FspI and AatII and separated on a denaturing polyacrylamide gel. Nascent strands generated by the leftward replication fork are indicated in brackets (lower panel). Extension products are indicated in the upper panel. Note that in pDPC^2xLead^ plasmids, both top and bottom strands are crosslinked to M.HpaII. Thus, extensions of both DNA strands appear with identical kinetics.

Since degradation of me-DPCs was completely blocked by SPRTN depletion, we asked whether overall replication and repair of me-DPCs is likewise dependent on SPRTN activity. To this end, we depleted SPRTN from extracts and monitored replication of pDPC^2xLead^ or pme-DPC^2xLead^. As seen before, in the context of an unmethylated DPC, SPRTN depletion only moderately delayed the conversion of gapped pDPC^2xLead^ to supercoiled repair products (Figure 5E, lanes 6-10). In contrast, SPRTN depletion caused a much greater stabilization of gapped pme-DPC^2xLead^ molecules (Figure 5E, lanes 16-20). This correlated with a defect in TLS, as seen by the accumulation of leading strands at the DPC (Figures 5F and 5G lower panel, lanes 13-16) and a corresponding delay in extension of leading strands past the lesion (Figure 5G upper panel, lanes 13-16). Strikingly, despite this defect in TLS, the upstream approach of leading strands to the lesion was largely unaffected by the combined inhibition of the protease pathways (Figure 5G, lower panel, lanes 13-16). Thus, in the absence of the proteasome pathway, SPRTN-dependent DPC proteolysis is exclusively required for efficient translesion DNA synthesis across the lesion site.

We next explored the importance of SPRTN’s protein-interacting regions for its role as a DPC protease. In addition to its SprT metalloprotease domain, SPRTN contains C-terminal p97 (SHP), PCNA (PIP), and ubiquitin (UBZ) interacting regions (Figure S5D). We generated recombinant SPRTN with mutated SHP, PIP, or UBZ domains (Figures S5D-F; (Davis et al., 2012; Mosbech et al., 2012)) and tested their activity in pme-DPC^2xLead^ replication. Whereas SPRTN depletion blocked degradation of me-DPCs during replication, re-addition of SPRTN-SHP* or SPRTN-PIP* reverted this effect (Figure 5D, lanes 8-9 and 10-11), suggesting that the p97 and PCNA binding interactions of SPRTN are not essential for its role as a DPC protease. In contrast, SPRTN-UBZ* failed to restore DPC proteolysis after SPRTN depletion (Figure 5D, lanes 12-13), suggesting that the ubiquitin binding function of SPRTN is important for DPC repair. The generation of supercoiled pme-DPC^2xLead^ repair products was likewise supported by SPRTN-SHP* and SPRTN-PIP* but not by SPRTN-UBZ* (Figure S5G). In summary, our data demonstrates that SPRTN acts on DPCs in the absence of DPC-ubiquitylation. Nevertheless, SPRTN activity is still dependent on its UBZ domains, suggesting that SPRTN interacts with a ubiquitylated protein other than the DPC at the lesion site.

### Single strand DNA triggers DPC ubiquitylation and degradation

We next addressed how SPRTN and proteasome activities are coupled to DNA replication. In one scenario, the replisome directly recruits or activates these proteases. Alternatively, DNA replication generates a structure that targets the proteases to DPCs. Consistent with the latter view, purified Wss1 and SPRTN are activated by single strand DNA (ssDNA) (Balakirev et al., 2015; Stingele et al., 2016). To examine this question in egg extracts, we tested whether ssDNA could trigger DPC degradation in the absence of the replisome. To this end, we generated a plasmid where M.HpaII is linked to one strand across from a 29 nt gap (pDPC^ssDNA^; see schematic in Figure S6A). We then monitored M.HpaII degradation on pDPC^ssDNA^ or pDPC in extracts that do not support MCM2-7 loading or replication initiation (non-licensing extracts). Strikingly, unlike pDPC, pDPC^ssDNA^ triggered rapid polyubiquitylation and degradation of M.HpaII (Figure 6A, lanes 5-9). As seen in the context of DNA replication, ubiquitylated M.HpaII was greatly stabilized by the combined inhibition of the proteasome and depletion of SPRTN (Figure 6B, lanes 10-12). Therefore, both SPRTN and the proteasome can degrade DPCs in the absence of the replisome when the lesion resides on ssDNA.

**Figure 6.**
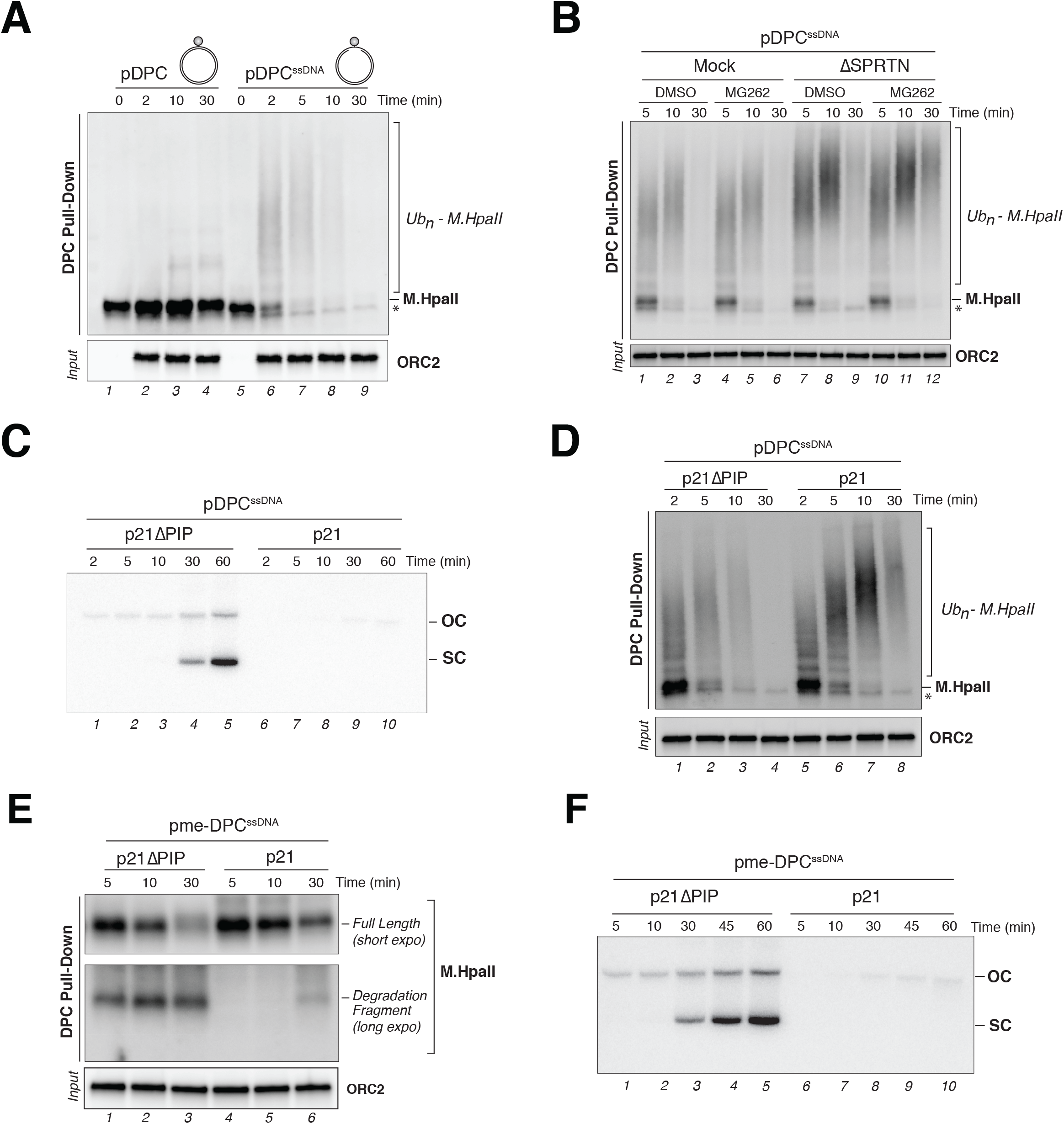
DPC ubiquitylation and degradation can occur in the absence of the replisome. (**A**) pDPC and pDPC^ssDNA^ were incubated in egg extracts that do not support MCM2-7 licensing (non-licensing). DPCs were recovered and monitored with a M.HpaII antibody as in Figure 1B. Note that time 0 were withdrawn prior to incubating plasmids in egg extracts explaining the absence of ORC2 input in lanes 1 and 5. (**B**) pDPC^ssDNA^ was incubated in mock-depleted and SPRTN-depleted non-licensing extracts. DMSO or 200 μM MG262 was supplemented to extracts where indicated. DPCs were recovered and monitored as in Figure 1B. (**C**) pDPC^ssDNA^ was incubated in non-licensing egg extracts in the presence of [α-^32^P]dATP. Extracts were supplemented with 500 μM of p21 peptide (p21) or p21 peptide with a mutated PIP box (p21ΔPIP) (Mattock et al., 2001). Samples were analyzed by agarose gel electrophoresis as in Figure 3B. (**D**) Samples from (C) were used to monitor DPC degradation as in Figure 1B. (**E**) pme-DPC^ssDNA^ was incubated in non-licensing egg extracts supplemented with 500 μM p21 or 500 μM p21ΔPIP peptide. DPC degradation was monitored like in Figure 1B. (**F**) Samples from (E) were supplemented with [α- ^32^P]dATP and analyzed by agarose gel electrophoresis as in Figure 3B. Note that SPRTN-dependent DPC degradation appears at 30 minutes (E, lane 6) at the same time that [α-^32^P]dATP incorporation is detected in lane 8.

When pDPC^ssDNA^ was incubated in non-licensing egg extracts, we detected low levels of DNA synthesis (Figure S6B). This synthesis reflects extension of the free 3’ end to the DPC, followed by translesion DNA synthesis past the lesion (Figure S6A). We asked whether this gap filling reaction is required to trigger DPC ubiquitylation and degradation. To this end, we pretreated egg extracts with a high concentration of p21 peptide, which contains a high affinity PCNA PIP box and therefore blocks interactions between PCNA and PIP box-containing proteins (Mattock et al., 2001) (Figure S6C). Addition of p21 peptide (p21), but not a mutant p21 peptide with a defective PIP box (p21ΔPIP), strongly inhibited gap filling (Figure 6C, lanes 6-10), presumably by blocking the PCNA-dependent recruitment of DNA polymerases. Strikingly, M.HpaII degradation was greatly delayed by p21, although M.HpaII ubiquitylation occurred normally (Figure 6D). Similar results were observed when gap filling was inhibited with a high concentration of aphidicolin, which blocks polymerase activity (Figures S6D and S6E). Thus, efficient DPC proteolysis but not DPC ubiquitylation requires polymerase extension to the lesion site.

Given that DPC degradation but not ubiquitylation was delayed by p21 peptide or aphidicolin, we hypothesized that DPC proteolysis by SPRTN but not the proteasome requires gap filling. Consistent with this model, inhibiting the proteasome in the presence of p21 led to a further accumulation of ubiquitylated M.HpaII compared to p21 alone (Figure S6F). To specifically monitor SPRTN activity, we examined a gapped substrate containing methylated M.HpaII, which cannot be acted on by the proteasome. As seen during DNA replication, methylated M.HpaII underwent ubiquitylation-independent degradation, giving rise to the SPRTN-dependent proteolytic fragment (Figures S6G, lanes 6-10). In this context, p21 significantly stabilized the methylated DPC, and dramatically delayed appearance of the proteolytic fragment (Figure 6E, lanes 4-6). The eventual appearance of the DPC fragment correlated with some residual gap filling (Figure 6F, lanes 6-10). In contrast, when p21 peptide was added after 3 minutes, when 3’ ends had already been extended (Figure S6A) but *prior* to the appearance of the SPRTN-dependent M.HpaII fragment, M.HpaII degradation occurred normally (Figure S6H), although TLS was impaired as indicated by defective supercoiling (Figure S6I). We conclude that strand extension close to the DPC is a prerequisite to trigger SPRTN-dependent DPC degradation. In contrast, a ssDNA gap appears to be sufficient to trigger DPC ubiquitylation and degradation via the proteasome.

### Polymerase extension controls SPRTN-dependent DPC degradation

Our experiments with the gapped substrate indicate that SPRTN-dependent DPC degradation is triggered by the extension of a DNA strand to the site of the DPC. To test this prediction in the context of a replication fork, a short peptide adduct was placed 16 nt upstream of the methylated DPC. The resulting pme-DPC^+peptide^ substrate was replicated in REV1-depleted egg extracts to cause permanent leading strand arrest at the peptide (Figure 7A). A matched pme-DPC substrate lacking the peptide served as a control. As seen in Figure 7B, after first pausing at the −30 to −40 positions due to CMG collision with the DPC, leading strands on pme-DPC^+peptide^ were extended but then permanently stalled at the upstream peptide adduct (lanes 5-8; −17, −16 positions). Under these conditions, methylated M.HpaII persisted, and the SPRTN-dependent proteolytic fragment never appeared (Figure 7C, lanes 5-7). In contrast, M.HpaII proteolysis proceeded normally on pme-DPC (Figure 7C), where leading strands were allowed to reach the DPC (Figure 7B). As an independent means of blocking strand extension, we added aphidicolin to extracts immediately after CMGs stalled at the DPC, and observed a similar inhibition of SPRTN-dependent DPC proteolysis (Figure S7A-D). Collectively, these results demonstrate that in the context of replication, DPC degradation by SPRTN is strictly dependent on polymerase extension to the lesion.

**Figure 7.**
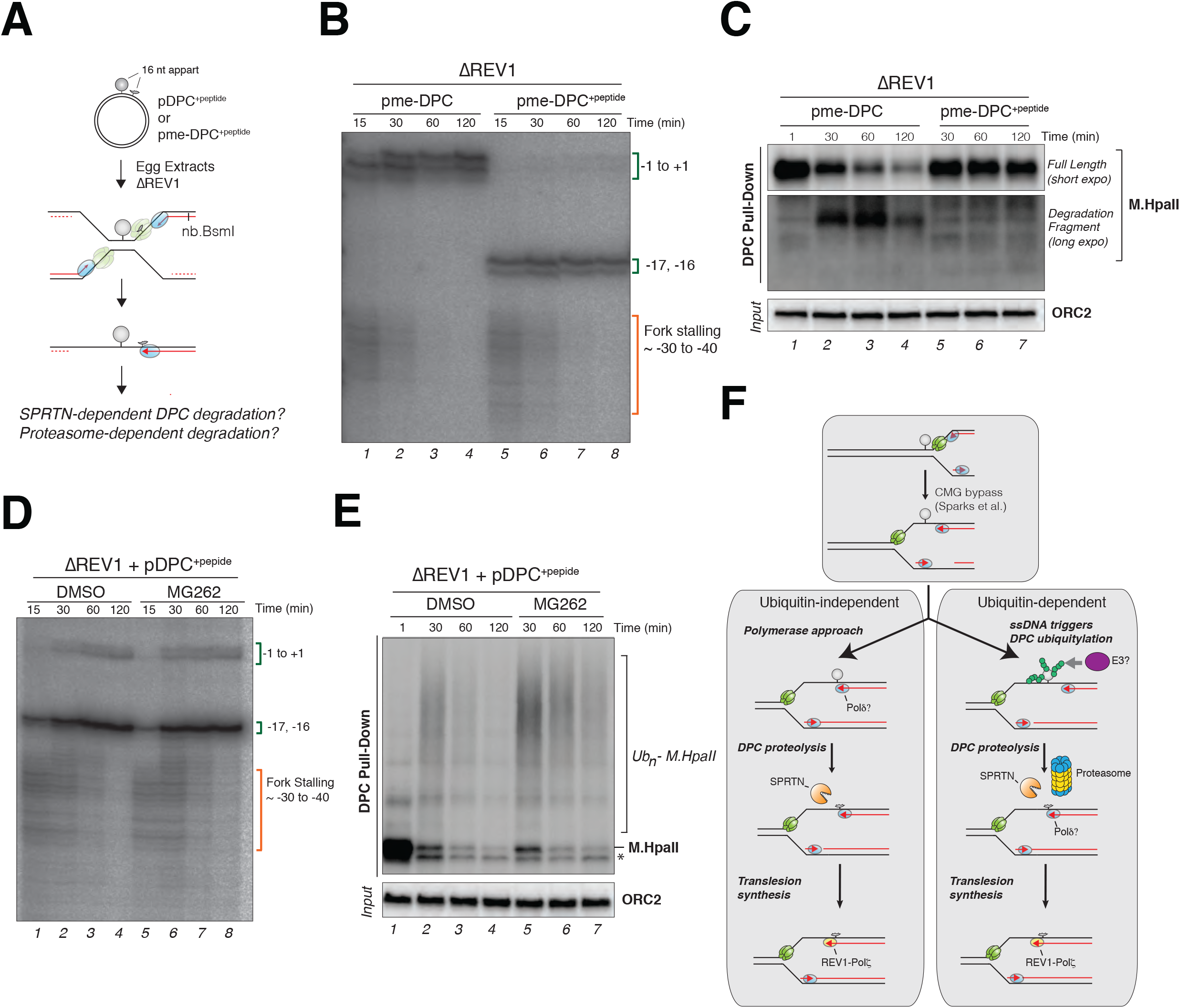
SPRTN-dependent DPC degradation requires polymerase extension to the lesion. (**A**) Depiction of pDPC+^peptide^ and pme-DPC+^peptide^ replication. (**B**) pme-DPC and pme-DPC+^peptide^ were replicated in REV1-depleted extracts in the presence of [α-^32^P]dATP. Samples were digested with nb.BsmI which specifically cuts the leftward leading strand as depicted in (A). Nascent leading strands were then separated on a polyacrylamide denaturing gel. Note that in the absence of REV1, nascent leading strands are unable to bypass the crosslink and permanently stall at −1, 0, +1 in pme-DPC, and −17,-16 in pme-DPC+^peptide^. CMG disappearance is monitored by the loss of the fork stalling signal (~ −30 to −40 positions) which occurs with identical kinetics in the pme-DPC and pme-DPC+^peptide^ samples. (**C**) Samples from (B) were used to monitor DPC degradation like in Figure 1B. Both short and long exposure of the M.HpaII blot are shown where indicated. Note the complete inhibition of the SPRTN-dependent degradation during replication of pme-DPC+^peptide^. (**D**) pDPC^+peptide^ was replicated in REV1 depleted extracts supplemented with either DMSO or 200 μM MG262 where indicated. Nascent leading strands were then separated on a polyacrylamide denaturing gel as in (B). (**E**) Samples from (D) were used to monitor DPC degradation as in Figure 1B. Note that MG262 treatment significantly stabilizes polyubiquitylated M.HpaII during replication of pDPC+^peptide^. (**F**) Model for replication-coupled DPC proteolysis in *Xenopus* egg extracts. DPC degradation can occur in the presence (Ubiquitin-dependent) or absence (Ubiquitin-independent) of DPC ubiquitylation. Black lines, parental DNA; red lines, nascent DNA, in green, CMG helicase; in blue, replicative polymerases; in yellow, TLS polymerase; in grey, DPC; in orange; SPRTN; in yellow and blue, the proteasome.

Having defined the requirements for SPRTN proteolysis, we repeated the same experiment with unmethylated M.HpaII, which should be degraded by the proteasome in the absence of polymerase extension. To this end, we replicated pDPC^+peptide^ in REV1-depleted extracts in the presence or absence of MG262. Unmethylated M.HpaII underwent rapid polyubiquitylation and degradation, although leading strands never reached the lesion due to the presence of the peptide adduct (Figures 7D, lane 1-4 and 7E, lanes 1-4). In this context, MG262 significantly stabilized polyubiquitylated M.HpaII (Figure 7E, lanes 5-7). Thus, DPC ubiquitylation and degradation by the proteasome do not require polymerase advancement to the lesion site. Based on experiments with pDPC^ssDNA^, we propose that ssDNA generated at stalled replication forks is sufficient to trigger DPC-ubiquitylation and activate the proteasome pathway.

## Discussion

We previously demonstrated that DPCs are degraded in a replication-dependent process, but how this occurs was unclear (Duxin et al., 2014). Using a newly developed PP-MS proteomic work flow, we identified SPRTN and the proteasome as two proteases recruited to a DPC lesion during DNA replication. We further demonstrate that SPRTN and the proteasome have partially overlapping functions in DPC proteolysis and are differentially activated by DNA replication. On one hand, SPRTN-dependent DPC degradation is tightly coupled to the extension of nascent DNA strands to the DPC which appears to be the underlying event that targets the protease (Figure 7F). In contrast, proteasome targeting to DPCs requires polyubiquitylation of the crosslinked protein, a process that is triggered by formation of single-stranded DNA near the lesion (Figure 7F). Our data demonstrate the existence of different mechanisms of S phase DPC repair, the implications of which we discuss below.

### Polymerase approach targets SPRTN

We demonstrate that following replisome-DPC encounter, SPRTN activity is dependent on the subsequent extension of a nascent DNA strand to within a few nucleotides of the DPC (Figure 7 AC), raising the possibility that the collision of a DNA polymerase with the DPC targets and activates SPRTN. Intriguingly, previous studies in mammalian cells reported direct interactions between SPRTN and POLD3, one of the accessory subunits of DNA Pol δ (Ghosal et al., 2012; Kim et al., 2013), suggesting that Pol δ might direct SPRTN to the DPC. To date, we have not been able to achieve sufficient depletion of Pol δ from egg extracts to prevent gap filling (data not shown), precluding a direct test of this model. Importantly, purified SPRTN is activated by DNA in vitro (Lopez-Mosqueda et al., 2016; Morocz et al., 2017; Stingele et al., 2016; Vaz et al., 2016), with ssDNA being particularly potent (Stingele et al., 2016). We therefore speculate that SPRTN activation requires the presence of a DNA polymerase on one side of the DPC and a short tract of ssDNA on the other side. This dual requirement would specifically target SPRTN to DPCs during replication and avoid the indiscriminate destruction of replisome or other chromatin components.

A model of SPRTN activation by polymerase-DPC collision has numerous implications and appealing features. First, because CMG blocks the ability of leading strands to reach a DPC on the leading strand template, our data imply that proteolysis by SPRTN can only occur if CMG is no longer present in front of the DPC (Figure 7F). Sparks et al. (submitted, see accompanying manuscript) show that the CMG helicase readily bypasses leading strand DPCs and that this process requires the helicase activity of RTEL1. Consistent with our model, in the absence of RTEL1-dependent DPC bypass, DPC proteolysis by SPRTN is suppressed. Second, SPRTN-dependent DPC proteolysis can likely be uncoupled from the replication fork. Supporting this idea, we show that SPRTN efficiently degrades a DPC attached to ssDNA in the absence of the replisome via a process that mimics post-replicative repair (Figure 6). By restricting SPRTN to act behind the replication fork, cells ensure that irreplaceable replisome factors such as CMG are not accidentally degraded during replication. Finally, SPRTN activation by polymerase-DPC collision suggests a common mechanism of DPC degradation on the leading and lagging strands: if Pol ε remains associated with CMG during DPC bypass, this would liberate the leading strand for Pol δ recruitment. In this way, both leading and lagging strand DPC proteolysis would be triggered by collision of Pol δ with the adduct.

Several reports originally described SPRTN as an important regulator of TLS in response to UV-induced DNA damage (Centore et al., 2012; Davis et al., 2012; Ghosal et al., 2012; Kim et al., 2013; Mosbech et al., 2012). Given the insight that SPRTN functions as a DPC protease, it is unclear whether SPRTN directly regulates TLS. We show that SPRTN stimulates TLS at a DPC but not a short peptide adduct, supporting the idea that SPRTN promotes efficient TLS indirectly via DPC proteolysis. In accordance with our results, experiments *in vitro* have shown that TLS polymerases can bypass short peptide DNA-adducts but not larger DPCs (Wickramaratne et al., 2016; Yamanaka et al., 2010; Yeo et al., 2014). Therefore, the increased mutagenesis observed in UV-irradiated cells lacking SPRTN (Kim et al., 2013) might reflect the accumulation of UV-induced DPCs that provoke more error-prone TLS than the corresponding peptide adduct.

SPRTN contains C-terminal domains that interact with p97, PCNA, and ubiquitin, but their importance for SPRTN activity is unclear. While some reports suggested that these domains are not essential (Maskey et al., 2014; Stingele et al., 2016), more recent evidence indicates that both the ubiquitin and PCNA binding interactions of SPRTN are important for its role as a DPC protease in human cells (Morocz et al., 2017). In *Xenopus* egg extracts, the ubiquitin binding domain of SPRTN is important for efficient DPC proteolysis, even in the absence of DPC ubiquitylation, suggesting that SPRTN interacts with another ubiquitylated protein near the lesion. Given our model that polymerase-DPC collision triggers SPRTN activity, a possible candidate is PCNA, which is ubiquitylated during post-replicative repair (Mailand et al., 2013). Consistent with this idea, RAD18, the E3 ligase that ubiquitylates PCNA, is epistatic to SPRTN for DPC repair (Morocz et al., 2017). Although the PCNA binding (PIP) motif was not required for SPRTN activity in egg extracts, this might reflect the presence of tandem UBZ domains in *Xenopus* SPRTN that could compensate for reduced PCNA binding.

### DPC ubiquitylation targets the proteasome

It was previously proposed that the proteasome can degrade DPCs (Baker et al., 2007; Lin et al., 2008; Mao et al., 2001; Quiñones et al., 2015; Reardon and Sancar, 2006), but direct evidence that this process can occur during DNA replication was lacking. Here, we show that DNA replication triggers rapid polyubiquitylation of a type I DPC, which is critical for its degradation by the proteasome. Importantly, DPC ubiquitylation occurs independently of the replisome or DNA synthesis, but requires ssDNA near the DPC. Thus, the E3 ligase that targets DPCs is likely activated by ssDNA. However, any protein residing on ssDNA should not be a target, as tracts of ssDNA are frequently generated in cells, e.g. during repair or helicase uncoupling at discrete DNA adducts. In some cases, as during nucleotide excision repair, ssDNA might never be accessible to the E3 ligase. In other cases, it might be too short-lived to support enough ubiquitylation to promote proteolysis. The discrimination between DPCs and non-covalent DNA binding proteins might be dictated not only by ubiquitylation but also by de-ubiquitylation of non-crosslinked proteins. Further studies are needed to understand how DPCs are recognized and targeted to the proteasome pathway.

### SPRTN and the proteasome are not redundant DPC proteases

How do our findings of two proteolytic pathways account for the defective replication fork progression observed in SPRTN-deficient cells (Lessel et al., 2014; Vaz et al., 2016)? We show above that whereas SPRTN depletion resulted in a TLS defect (Figure 3), proteasome inhibition did not (Figure 4), indicating that these proteolytic functions are not completely redundant. We reason that SPRTN is able to degrade DPCs to peptide adducts that are sufficiently small for efficient TLS. Indeed, the protease active site of Wss1 is highly solvent exposed, suggesting it should be able to cleave DPCs close to the DNA attachment site (Stingele et al. 2016). In contrast, the active sites of the proteasome are buried inside the 20S core particle. DPC proteolysis by the proteasome therefore requires threading of the unfolded DPC through the cylindrical 20S particle, which would likely be interrupted upon encounter with the attached DNA, resulting in a larger peptide adduct. Thus, when DPCs are channeled into the proteasomal degradation pathway, SPRTN may still be required in a second proteolytic step to degrade the peptide adduct down to a few amino acids (Figure 7F). Our findings predict that in SPRTN-deficient cells, CMG becomes uncoupled from the leading strand due to defective TLS. In bacteria, helicase uncoupling greatly slows the rate of DNA unwinding (Kim et al., 1996). Therefore, we speculate that defective fork progression in SPRTN-deficient cells reflects slow unwinding by uncoupled CMG.

### Relevance of the proteasome pathway

DPCs are expected to exhibit great variability in size, structure, and attachment chemistry. While agents such as formaldehyde crosslink proteins to duplex DNA, abortive reactions by topoisomerase form DPCs that are flanked by a DNA break (Barker et al., 2005; Ide et al., 2011; Stingele et al., 2017). Hence, while both SPRTN and the proteasome readily degrade M.HpaII, it is conceivable that other crosslinked proteins are preferentially processed by one or the other protease. For example, SPRTN-mediated DPC proteolysis is expected to be particularly critical for DPCs that lack available lysines and cannot be ubiquitylated (Figure 5). In contrast, proteasome-dependent DPC degradation might be essential for very large DPCs that cannot be bypassed by the replication fork and require “pre-trimming” by the proteasome. Alternatively, the essential role of the proteasome pathway may be independent of DNA replication. Instead, the proteasome might be critical to remove DPCs flanked by DNA breaks prior to DNA replication as in the case of topoisomerase I (Desai et al., 1997). By employing at least two DPC proteases with orthogonal mechanisms and triggers, cells are equipped with a versatile system that can efficiently degrade a wide variety of DPCs.

## Experimental Procedures

### Xenopus Egg Extracts and DNA Replication

Preparation of *Xenopus* egg extracts was performed as described previously (Lebofsky et al., 2009). For DNA replication, plasmids were first incubated in a high-speed supernatant (HSS) of egg cytoplasm (final concentration of 7.5-15 ng DNA/μL HSS) for 20-30 min at room temperature to license the DNA, followed by the addition of two volumes of nucleoplasmic egg extract (NPE) to initiate replication. Where indicated, HSS was supplemented with Geminin at a final concentration of 10 μM and incubated for 10 min at room temperature prior to addition of plasmid DNA. For replication in the presence of LacI, plasmid DNA (75 ng/uL) was incubated with an equal volume of 12 μM LacI for 1 hr prior to HSS addition (Duxin et al., 2014). For UbVS treatment, NPE was supplemented with 22.5 μM ubiquitin vinyl sulfone (UbVS) (Boston Biochem) and incubated for 15 min prior to mixing with HSS (15 μM final concentration). Where indicated, recombinant ubiquitin or FLAG-ubiquitin (Boston Biochem) were added to NPE at a concentration of 120 μM (80 μM final concentration). For SPRTN depletion-rescue experiments, NPE was supplemented with 30 nM recombinant wild type or mutant *Xenopus* SPRTN. For SPRTN-EQ dominant negative experiments, recombinant *Xenopus* SPRTN-EQ was added to undepleted NPE at a concentration of 400 nM. To block de novo SUMOylation, dnUBC9 was added to extracts to a final concentration of 10 μM (Azuma et al., 2003). Where indicated, proteasome activity was inhibited via the addition of 200 μM MG262 (Boston Biochem) to extracts (final concentration). For DNA labeling, reactions were supplemented with [α-^32^P]dATP. To analyze plasmid replication intermediates, 1 uL of each reaction was added to 10 μL of replication stop solution A (5% SDS, 80 mM Tris pH 8.0, 0.13% phosphoric acid, 10% Ficoll) supplemented with 1 μL of Proteinase K (20 mg/ml) (Roche). Samples were incubated for 1 hr at 37°C prior to separation by 0.9% native agarose gel electrophoresis and visualization using a phosphorimager (Lebofsky et al., 2009). For analysis of nascent leading strand products, 3-4 μL of each replication reaction was added to 10 volumes of 50 mM Tris pH 7.5, 0.5% SDS, 25 mM EDTA, and replication intermediates were purified as previously described (Räschle et al., 2008). For incubation in non-licensing extracts (Figures S1C, 6A-F and S6A-I), one volume of HSS and two volumes of NPE were premixed prior to the addition of plasmid DNA (final concentration of 15 ng/μL). Where indicated, p21 peptide (Ac-CKRRQTSMTDFYHSKRRAIAS-amide) or p21ΔPIP peptide (Ac-CKRRATSATDAAHSKRRAIAS-amide) was added to HSS/NPE mix to a final concentration of 500 μM (Mattock et al., 2001). All experiments were performed at least in duplicate and a representative experiment is shown.

#### Preparation of DNA constructs

To generate pDPC we first created pJLS2 by replacing the AatII-BsmBI fragment from pJD2 (Duxin et al., 2014) with the following sequence 5’-GGGAGCTGAATGCCGCGCGAATAATGGTTTCTTAGACGT-3’ which contains a nb.BsmI site. To generate pDPC^2xLead^, the SacI-BssHII fragment from pJLS2 was replaced with the following sequence: 5’-CATCCACTAGCCAATTTATGCTGAGGTACCGGATTGAGTAGCTACCGGATGCTGAGGGGATCCACTAGCCAATTTATCATGG-3’. pJLS2 or pJLS3 were nicked with nt.BbvcI and ligated with the following oligo containing a fluorinated cytosine: 5’-TCAGCATCCGGTAGCTACTCAATC[C5-Fluro dC]GGTACC-3’ and subsequently crosslinked to M.HpaII-His_6_ to generate pDPC or pDPC^2xLead^, respectively, as previously described (Duxin et al., 2014). To generate pDPC^PK^, pDPC was treated with Proteinase K (37°C overnight in presence of 0.5% SDS) to reduce the DPC to a 4 amino acids peptide adduct. The plasmid was subsequently recovered by phenol/chloroform extraction. To generate pDPC^+peptide^ the ApoI-NdeI fragment of pJLS2 was replaced with the following sequence: 5’-AATTCCTCAGCATCCGGTTCGAACTCAATAGCTTACCTCAGCCA-3’, generating pNBL104. pNBL104 was nicked with Nt.BbvCI and ligated with the following oligo containing both a fluorinated AluI site and a fluorinated M.HpaII site: 5’-TCAGCATC[C5-FlurodC]GGTTCGAACTCAATAG[C5-FlurodC]TTACC-3’. AluI Methyltransferase (New England BioLabs) was first crosslinked to the plasmid, degraded with Proteinase K (37°C overnight in presence of 0.5% SDS) and the plasmid was recovered by phenol/chloroform extraction. The peptide-containing plasmid was then crosslinked to M.HpaII-His_6_ as described above. To generate pDPC^ssDNA^, pJLS2 was nicked with nb.BbvCI and ligated with the following fluorinated oligo: 5’-TGAGGTAC[C5-FlurodC]GGATTGAGTAGCTACCGGATGC-3’. The dFdC-containing plasmid was cut with nt.BbvCI and the resulting 31bp fragment was melted off and captured by annealing to an excess complimentary oligo 5’-GGTACCGGATTGAGTAGCTACCGGATGCTGA-3’. Excess oligos were then degraded by Exonuclease I (New England BioLabs) treatment. The gapped plasmid was then recovered by phenol/chloroform extraction and crosslinked to M.HpaII-His_6_ as described above.

#### Antibodies and Immunodepletion

The following antibodies used were described previously: REV1 (Budzowska et al., 2015), ORC2 (Fang and Newport, 1993), and CDT1 (Arias and Walter, 2005). M.HpaII antibody was raised against full length M.HpaII-His6 expressed and purified from bacteria under denaturing conditions (Pocono Rabbit Farm & Laboratory). PSMA1, PSMA3, SPRTN, and MCM6 antibodies were raised by New England Peptide by immunizing rabbits with Ac-CAEEPVEKQEEPMEH-OH, Ac-CKYAKESLEEEDDSDDDNM-OH, Aoa-DVLQDKINDHLDTCLQNCNT-OH, and Ac-CLVVNPNYMLED-OH, respectively. SPRTN-F antibody was raised against a fragment of *Xenopus laevis* SPRTN encompassing amino acids 302-528 which was tagged on N-terminus with His_6_. The protein fragment was purified from bacteria under denaturing conditions and the antibody was raised by Pocono Rabbit Farm & Laboratory. Western blotting analysis for H3 was carried with commercial antibody from Cell Signaling (Cat #9715S).

To immunodeplete SPRTN from *Xenopus* egg extracts, one volume of Protein A Sepharose Fast Flow (PAS) (GE Health Care) was mixed with either 4 volumes of affinity purified SPRTN peptide antibody (1 mg/mL) or 1 volume of α-SPRTN-F serum and incubated overnight at 4°C. The beads were then washed twice with 500 μL PBS, once with ELB (10 mM HEPES pH 7.7, 50 mM KCl, 2.5 mM MgCl_2_, and 250 mM sucrose), three times with ELB supplemented with 0.5 M NaCl, and twice with ELB. One volume of precleared HSS or NPE was then depleted by mixing with 0.2 volumes of antibody-bound beads then incubating at room temperature for 20 min. The depletion procedure was repeated once. To immunodeplete PSMA1, one volume of PAS beads was mixed with 10 volumes of affinity purified PSMA1 peptide antibody (1 mg/mL). The beads were washed as described above, and one volume of precleared HSS or NPE was then depleted by mixing with 0.2 volumes of antibody-bound beads and then incubating at room temperature for 20 min. The depletion procedure was repeated three times for HSS and twice for NPE. For SPRTN and PSMA1 combined depletion, one volume of PAS beads was mixed with 4 volumes of affinity purified SPRTN peptide antibody and 10 volumes of affinity purified PSMA1 peptide antibody. The beads were washed and depletion was performed as described for PSMA1 immunodepletion. The immunodepletion of REV1 was performed as previously described (Budzowska et al., 2015).

#### Nascent-Strand Analysis

Nascent strand analysis was performed as previously described (Räschle et al., 2008). Briefly, purified DNA was digested with the indicated restriction enzymes followed by addition of 0.5 volumes of Gel Loading Dye II (Denaturing PAGE) (Life Technologies). DNA fragments were subsequently separated on 5% or 7% denaturing polyacrylamide gels, transferred to filter paper, dried, and visualized using a phosphorimager. The images shown in the lower panels of Figures 3D, S3D, S3F, 5G, 7B, 7D and S7B, were processed using logarithmic transformation in ImageJ to allow better visualization of fork stalling signal (~ −30 to −40 positions).

Reference oligo used in Figure S3G: 5’-CATTCAGCTCCCGGAGACGGTCACAGCTTGTCTGTAAGCGGATGCCGGGAGCAGACAAGCCCGTCAGGGCGCGTCAGCGGGTGTGGCGGGTGTCGGGGCTGGCTTAACTATGCGGCATCAGAGCAGATTGTACTGAGAGTGCACCATATGGCTGAGGTACCG-3’.

Primer used for dideoxy-sequencing ladder in Figure S3G: 5’-CAT TCA GCT CCC GGA GAC GGT C – 3’.

#### Protein Expression and Purification

M.HpaII-His_6_ and LacI-biotin were expressed and purified as previously described (Duxin et al., 2014). To generate lysine-methylated M.HpaII, purified M.HpaII-His_6_ was first denatured by dialyzing against 20 mM HEPES pH 7.5, 100 mM KCl, 6M Guanidine HCl, 10% glycerol. Denatured M.HpaII protein was then methylated using Reductive Alkylation Kit (Hampton Research) via the addition of dimethylamine borane and formaldehyde according to the manufacturer’s protocols. The methylation reaction was stopped by addition of 100 mM Tris pH 7.5 and 5 mM DTT (final concentrations). Methylated M.HpaII was then renatured by sequentially dialyzing against Renaturing Buffer (20 mM Tris pH 8.5, 100mM KCl, 1mM DTT, 10% glycerol) supplemented with 4, 2, and 0M Guanidine HCl for 1 hr each at 4°C. The renatured protein was then dialyzed against storage buffer (20 mM Tris pH 8.5, 100 mM KCl, 1 mM DTT, 30% glycerol) and stored at −80°C.

*Xenopus* SPRTN with an N-terminal FLAG tag was cloned into pFastBac1 (Thermo Fisher Scientific) using primers A and B. SPRTN mutations were introduced via Quikchange mutagenesis and confirmed by Sanger sequencing. SPRTN Baculoviruses were prepared using the Bac-to-Bac system (Thermo Fisher Scientific) according to the manufacturer’s protocols. SPRTN was expressed in 250 mL suspension cultures of Sf9 insect cells (Thermo Fisher Scientific) by infection with SPRTN baculovirus for 48 hr. Sf9 cells were subsequently collected via centrifugation and resuspended in Lysis Buffer (50 mM Tris pH 7.5, 500 mM NaCl, 10% Glycerol, 1X Roche EDTA-free Complete protease inhibitor cocktail, 0.5 mM PMSF, 0.2% Triton X-100). To lyse cells, the suspension was subjected to three freeze/thaw cycles, passed through a 21g needle, and then sonicated. The cell lysate was spun at 25000 rpm in a Beckman SW41 rotor for 1hr. The soluble fraction was collected and then incubated with 200 μL anti-FLAG M2 affinity resin (Sigma) for 90min at 4°C. The resin was then washed once with 10 mL Lysis Buffer, twice with Wash Buffer (50 mM Tris pH 7.5, 500 mM NaCl, 10% Glycerol, 0.2% Triton X-100), and three times with Buffer A (50 mM Tris pH 7.5, 500 mM NaCl, 10% Glycerol). FLAG-SPRTN was eluted with Buffer A supplemented with 100 μg/mL 3xFLAG peptide (Sigma). Elution fractions containing FLAG-SPRTN protein were pooled and dialyzed against 20 mM Tris pH 7.5, 300 mM NaCl, 10% Glycerol, 1mM DTT at 4°C for 12 hr and then dialyzed against Storage Buffer (20 mM Tris pH 7.5, 150 mM NaCl, 10% Glycerol, 1mM DTT) at 4°C for 3 hr. Aliquots of FLAG-SPRTN were then stored at −80°C. Primer A: 5’ – GAT CGG ATC CAT GGA CTA CAA AGA CGA TGA CGA CAA GGG TGA TAT GCA GAT GTC GGT AG − 3’ Primer B: 5’-GAT CCT CGA GTT ATT ATG TAT TGC AGT TTT GTA AGC AGG TGT CTA AAT G −3’

#### Plasmid Pull-Downs

Plasmid pull-down assays were performed as previously described (Budzowska et al., 2015). Proteins associated with the chromatin fraction were visualized by Western blotting with the indicated antibodies.

#### DPC Pull-Downs

We developed a modified plasmid pull-down protocol to specifically isolate M.HpaII DPCs from extracts (Figure 1A). Streptavidin-coupled magnetic beads (Dynabeads M-280, Invitrogen; 5uL per pull-down) were washed twice with 50 mM Tris pH 7.5, 150mM NaCl, 1mM EDTA pH 8, 0.02% Tween-20. Biotinylated LacI was added to the beads (1 pmol per 5 μL of beads) and incubated at room temperature for 40 min. The beads were then washed four times with DPC pull-down buffer (20 mM Tris pH 7.5, 150 mM NaCl, 2 mM EDTA pH 8, 0.5% IPEGAL-CA630) and then stored in the same buffer on ice until needed. At the indicated times during DNA replication or gap filling, equal volumes (2-10 μL) of reaction were withdrawn and stopped in 300 μL of DPC pull-down buffer on ice. After all of the time points were taken, 5 μL of LacI-coated streptavidin Dynabeads were added to each sample and allowed to bind for 30-60 min at 4°C rotating. 20 μL of pull-down supernatant was reserved in 20 μL of 2X Laemmli sample buffer for input. The beads were subsequently washed four times with DPC pull-down buffer and then twice with Benzonase buffer (20mM Tris pH 7.5, 150mM NaCl, 2mM MgCl_2_, 0.02% Tween-20) before being resuspended in 15 μL Benzonase buffer containing 1 μL Benzonase (Novagen). Samples were incubated for 1hr at 37°C to allow for DNA digestion and DPC elution, after which the beads were pelleted and the supernatant M.HpaII eluate was mixed with 2X Laemmli sample buffer for subsequent Western blotting analysis.

For FLAG immunoprecipitation analysis of isolated DPCs (Figure 1D), the M.HpaII eluate resulting from Benzonase treatment was instead diluted to 300 μL in Benzonase buffer. FLAG M2 magnetic beads (Invitrogen; 5 μL per pull-down) were added to each sample and allowed to bind for 60 min at 4°C rotating. The beads were subsequently washed four times with Benzonase buffer. To elute precipitated proteins, the beads were then resuspended in 0.1M Glycine pH 3 and incubated with gentle shaking for 10 min at room temperature. After pelleting the beads, the supernatant eluate was subsequently neutralized with 10mM Tris pH 11 and mixed with 2X Laemmli buffer.

To monitor M.HpaII degradation pDPC^Lead^ or pDPC^Lag^ plasmids were pre-bound with purified LacI (untagged) for 60 min at RT as previously described (Duxin et al., 2014). Pre-bound plasmids were replicated at 5 ng/uL final concentration in HSS/NPE, and reactions stopped in DPC pull-down buffer. DPC plasmids were pulled down washed and benzonase treated as described above. After elution with benzonase, the eluates were incubated with His-tag dynabeads (Life Technologies) in HIS wash buffer (50 mM sodium phosphate buffer, pH8, 150 mM NaCl, 0.02% Tween-20) for 10 minutes at 4 C. This step was added to avoid cross reactivity between free LacI and the M.HpaII antibody. Beads were washed three times in HIS wash buffer and eluted in HIS elution buffer (300 mM imidazole, 50 mM sodium phosphate buffer, pH8, 300 mM NaCl, 0.01% Tween20) shaking at RT for 5 min. The supernatant M.HpaII-His_6_ eluate was mixed with 2X Laemmli sample buffer for subsequent Western blotting analysis.

#### Plasmid Pull-down Mass Spectrometry (PP-MS)

Plasmid DNA was replicated in egg extracts at 5 ng/uL (final concentration). At the indicated time points, 8 μL of the reaction were withdrawn and plasmids and associated proteins were recovered by plasmid pull down using LacI coated beads (Budzowska et al., 2015). After 30 min incubation at 4°C, samples were washed twice in 10 mM HEPES pH 7.7, 50 mM KCl, 2.5 mM MgCl_2_, 0.03% Tween 20, and once in 10 mM HEPES pH 7.7, 50 mM KCl, 2.5 mM MgCl_2_. Samples were washed one additional time in 50 μL of 10 mM HEPES pH 7.7, 50 mM KCl, 2.5 mM MgCl_2_ and transferred to a new tube to remove residual detergent. Beads were dried out and resuspended in 50 μL denaturation buffer (8 M Urea, 100 mM Tris pH 8.0). Cysteines were reduced (1 mM DTT, 15 minutes at RT) and alkylated (5 mM iodoacetamide, 45 min at RT). Proteins were digested and eluted from beads with 1.5 μg LysC (Sigma) for 2.5 hr at RT. Eluted samples were transferred to a new tube and diluted 1:4 with ABC (50 mM ammonium bicarbonate). 2.5 μg trypsin was added and incubated for 16 hours at 30°C. NaCl was added to 400 mM final concentration, and peptides were acidified and purified by stage tipping on C18 material. Samples were analyzed on a Q Exactive HF Orbitrap mass spectrometer (Thermo Scientific) and quantified by the label free algorithm implemented in the MaxQuant software, as previously described (Räschle et al., 2015). MS experiments were carried out in quadruplicates. A fifth replicate was used to isolate the DNA repair intermediates shown in Figure 2A.

## Author Contributions

A.G, N.B.L and I.G performed the experiments. J.P.D and J.L.S prepared the M.S samples which were run and analyzed by M.R and M.M. J.C.W helped design and analyse experiments at early stages of the project. A.G, N.B.L and J.P.D designed and analyzed the experiments and prepared the manuscript.

## Acknowledgments

We thank Peter Burgers, Jiri Lukas, Niels Mailand and members of the Walter and Duxin laboratories for feedback on the manuscript. We thank Yoshiaki Azuma for dnUBC9 protein and expression construct. J.P.D’s lab is supported by grants from the Novo Nordisk Foundation (NNF14CC0001) and the European Research Council (ERC) under the European Union’s Horizon 2020 research and innovation programme (grant agreement No 715975). MR was supported by the Center for Integrated Protein Research Munich (CIPSM). J.C.W. was supported by NIH grant HL98316. J.C.W. is an investigator of the Howard Hughes Medical Institute.

---

**Figure S1.**
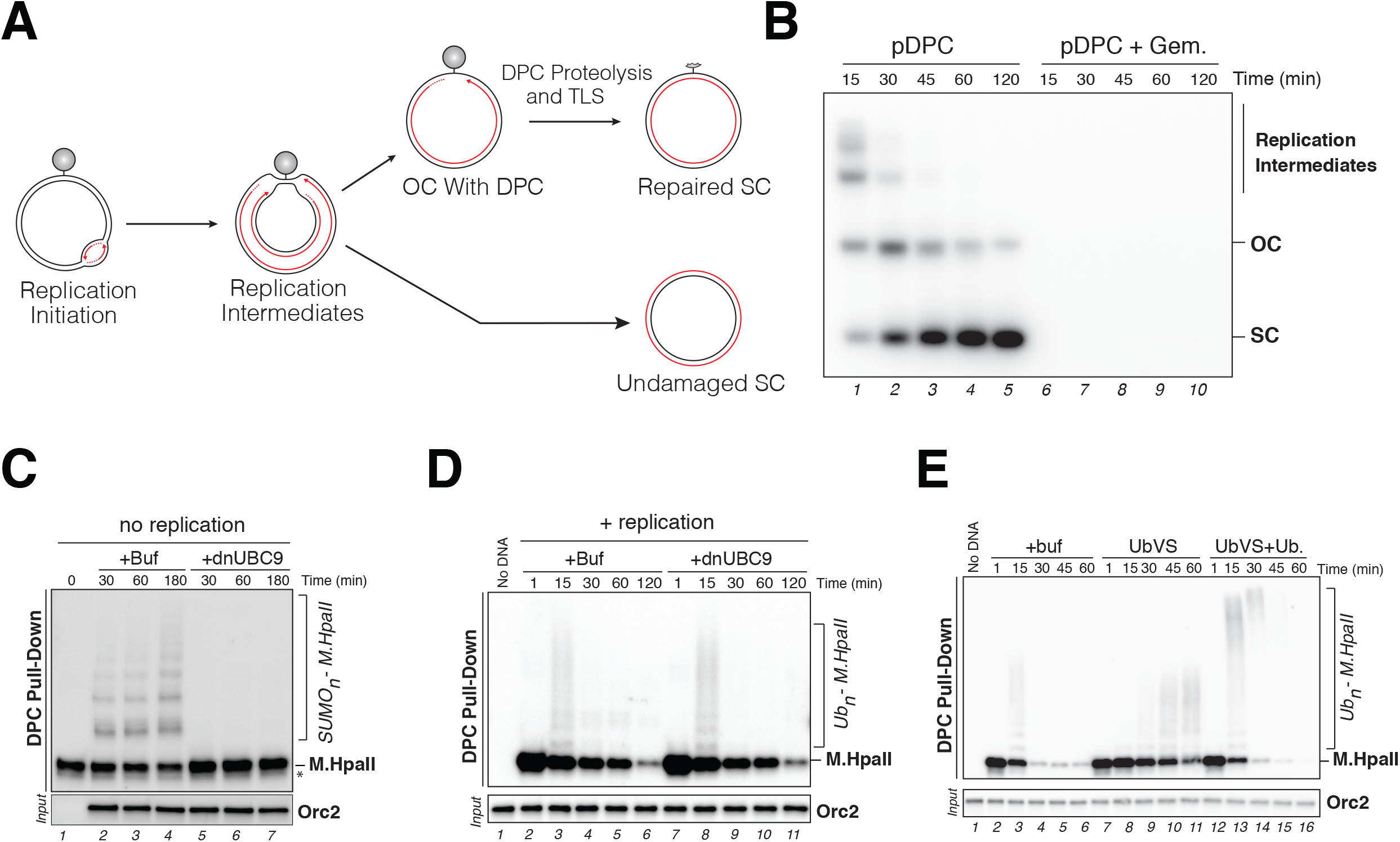
(**A**) Schematic illustrating replication intermediates generated during replication of pDPC (Duxin et al., 2014). (**B**) pDPC was replicated in egg extracts in the presence of [α-^32^P]dATP. Geminin (+Gem.) was supplemented where indicated to block DNA replication (Tada et al., 2001; Wohlschlegel et al., 2000). Samples were analyzed by agarose gel electrophoresis. OC, open circular; SC, supercoiled. (**C**) pDPC^2xLead^ was incubated in egg extracts that do not support CMG licensing and replication initiation. Recombinant UBC9 dominant-negative (+dnUBC9) was added where indicated to block SUMOylation (Azuma et al., 2003). DPCs were recovered and monitored as in Figure 1B. (**D**) pDPC^2xLead^ was replicated in egg extracts supplemented with buffer (+buf) or UBC9 dominant-negative (+dnUBC9). DPCs were recovered and monitored as in Figure 1B. Note that DPCs are polyubiquitylated and degraded with similar kinetics in the presence or absence of dnUBC9. (**E**) pDPC^2xLead^ was replicated in egg extracts in the presence of buffer (+buf), 15 μM of ubiquitin-vinyl-sulfone (UbVS), or 15 μM of UbVS and 80 μM of recombinant ubiquitin. DPCs were recovered and monitored as in Figure 1B.

**Figure S2.**
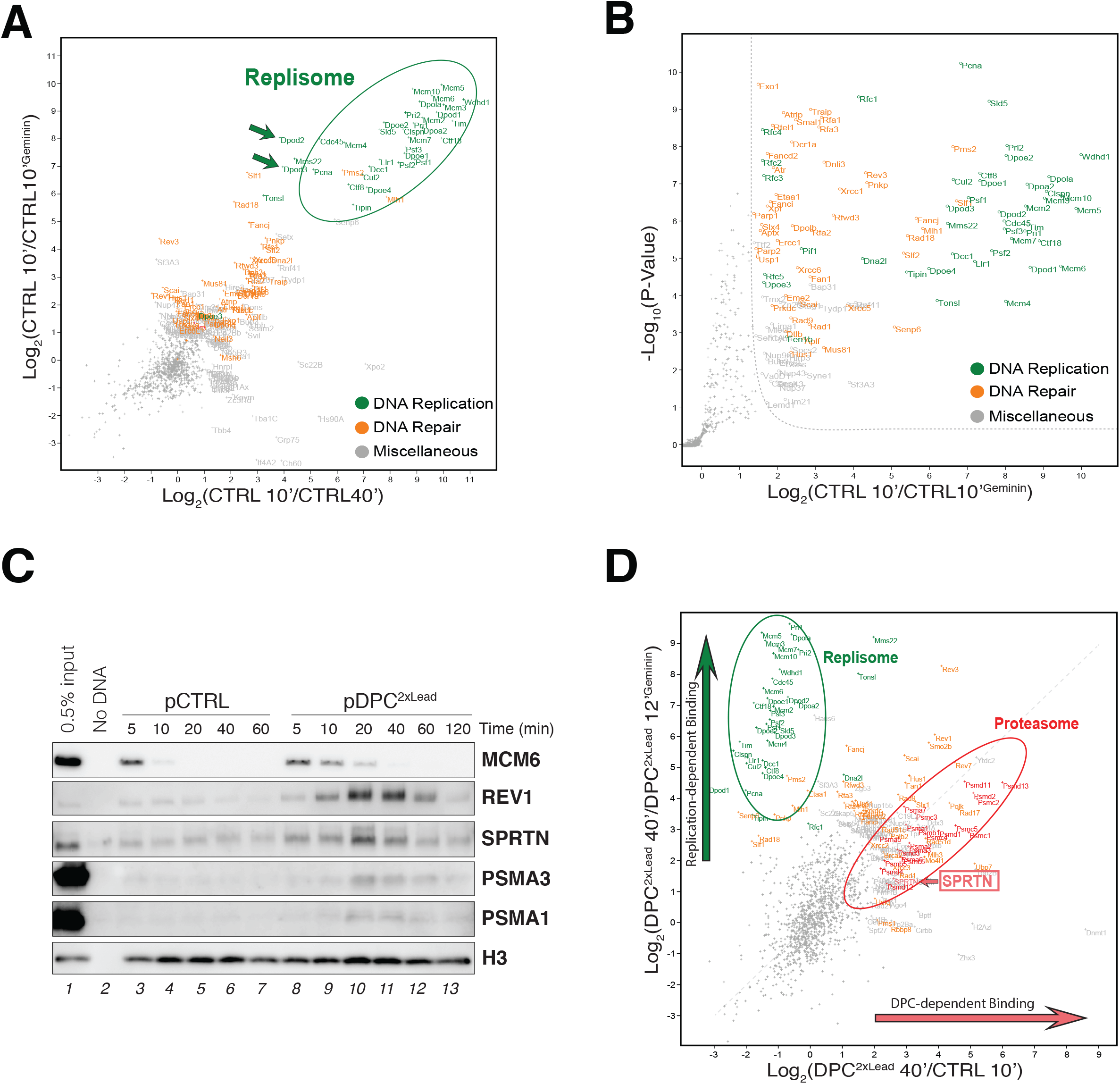
(**A**) Analysis of protein recruitment to pCTRL at 10 min compared to pCTRL at 40 min (x axis) and pCTRL at 10 min compared to pCTRL + Geminin at 10 min (y axis). The plot shows the mean difference of the protein intensity of 4 biochemical replicates for each of the conditions indicated. (**B**) Analysis of protein recruitment to pCTRL compared to pCTRL + Geminin at 10 min. The volcano plot shows the mean difference of the protein intensity plotted against the *P* value of 4 biochemical replicates. (**C**) pDPC^2xLead^ and pCTRL were replicated in egg extracts and recovered at the indicated time point as depicted in Figure 2A (Budzowska et al., 2015). Samples were blotted with the indicated antibodies. (**D**) Analysis of protein recruitment to pDPC^2xLead^ at 40 min compared to pCTRL at 10 min (x axis) and pDPC^2xLead^ at 40 min compared to pDPC^2xLead^ + Geminin at 12 min (y axis). The plot shows the mean difference of the protein intensity of 4 biochemical replicates for each of the conditions indicated.

**Figure S3.**
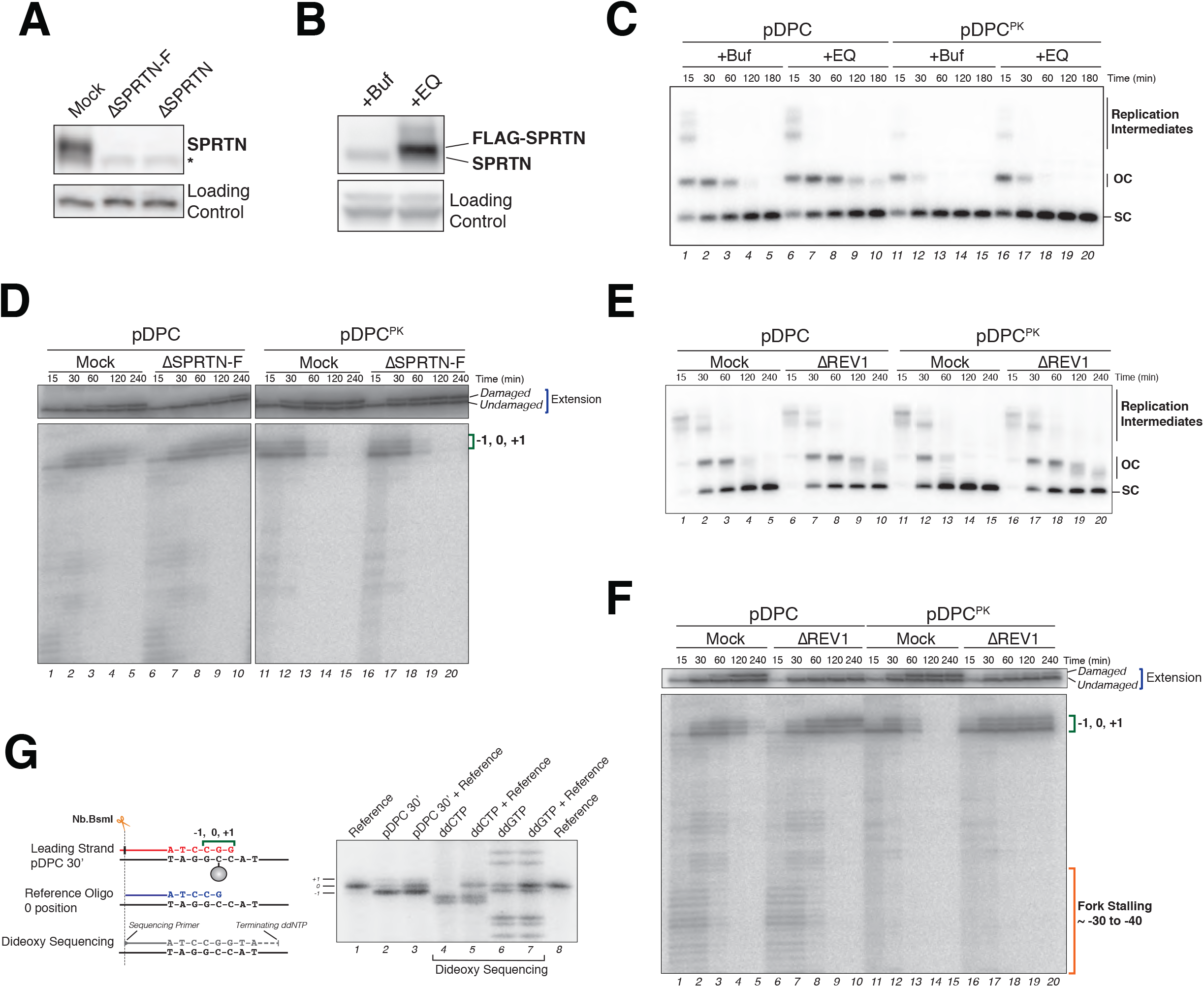
(**A**) Mock-depleted or SPRTN-depleted extracts were blotted against SPRTN or MCM6 (loading control). Two different antibodies generated against SPRTN (SPRTN-F and SPRTN) deplete the protein with similar efficiency. (**B**) Egg extracts were supplemented with buffer or recombinant SPRTN catalytically inactive (+EQ) and blotted with SPRTN or MCM6 (loading control) antibody. (**C**) Extracts from (B) were used to replicate pDPC or pDPC^PK^. Replication samples were analyzed by agarose gel electrophoresis as in Figure 3B. (**D**) Mock-depleted and SPRTN-depleted extracts were used to replicate pDPC. Samples were digested with NcoI and AatII and separated on a denaturing polyacrylamide gel. (**E**) Mock-depleted and REV1-depleted extracts were used to replicate pDPC and pDPC^PK^. Samples were analyzed by agarose gel electrophoresis as in Figure 3B. Note the accumulation of open circular molecules (OC) in the absence of REV1 indicative of TLS defect. OC molecules that accumulate decline in size over time (lanes 17-20) due to 5’ to 3’ end DNA resection activity present in egg extracts (Duxin et al., 2014). (**F**) Samples from (E) were digested with NcoI and AatII and separated on a polyacrylamide denaturing gel as in (D). (**G**) Nascent strand intermediates stall at −1, 0 and +1 and not at 0, +1 and +2 as previously reported (Duxin et al., 2014). 30 min samples of pDPC were analyzed by denaturing polyacrylamide gel electrophoresis following nb.BsmI digest alongside a reference oligo and a sequencing ladder generated with ddCTP and ddGTP. In lane 3, the 30 min sample of pDPC was premixed with the reference oligo before electrophoresis. In lanes 5 and 7, the reference oligo was premixed with the cytosine (ddCTP) and guanine (ddGTP) samples of the sequencing ladder, respectively. Note that the middle band of the three discrete bands generated during replication of pDPC aligns with the reference oligo (0 position). Note also that the sequencing ladder alignment with the reference oligo is shifted 1 nucleotide (lanes 4-8). This is likely caused by a faster migration of the sequencing ladder induced by the terminal dideoxynucleotide. In Duxin et al. 2014, the nascent strand stalling positions were determined by comparing pDPC to a sequencing ladder generated by dideoxy sequencing causing the incorrect assignment of the stalling intermediates.

**Figure S4.**
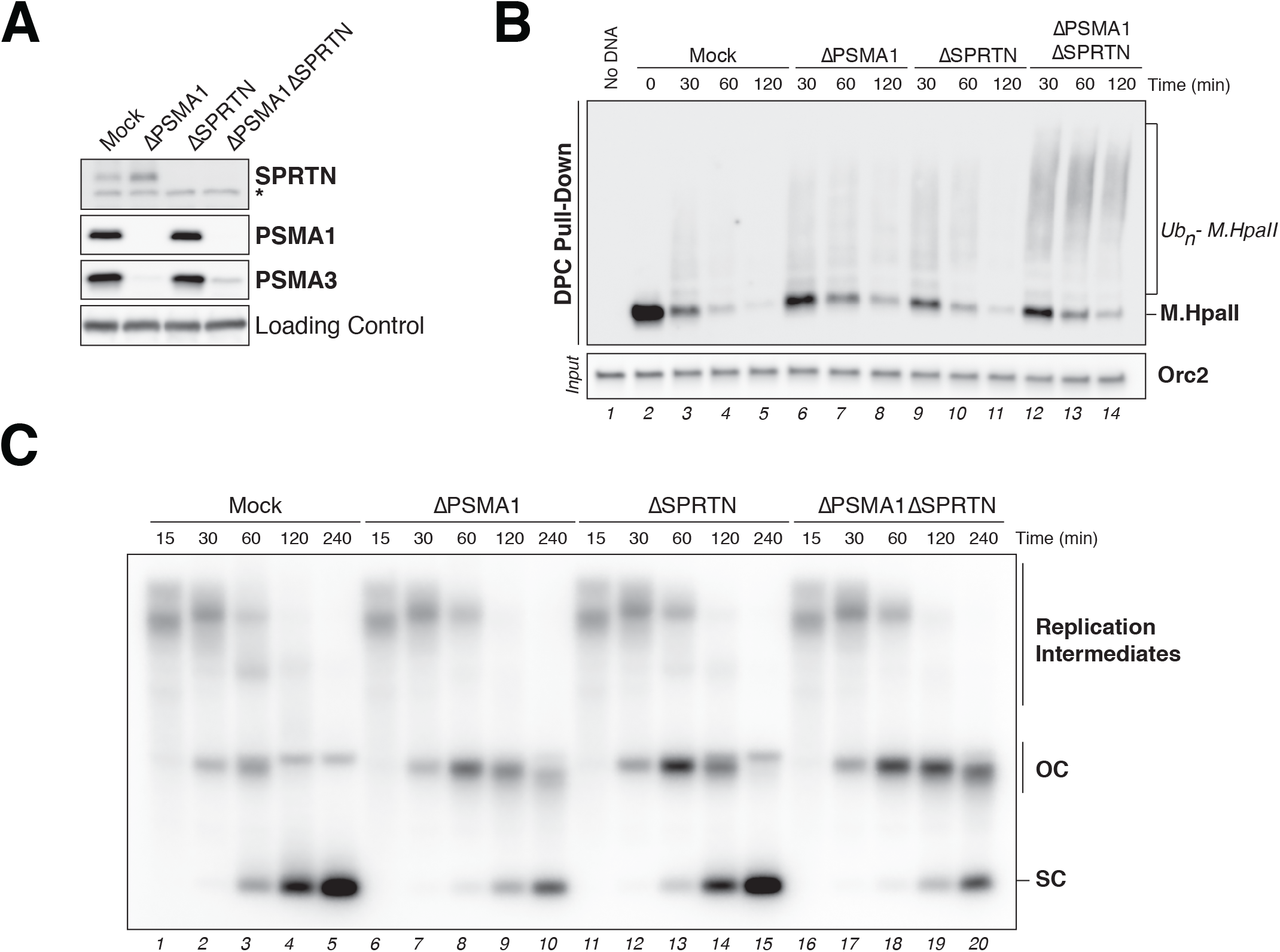
(**A**) Mock-depleted, PSMA1-depleted, SPRTN-depleted or PSMA1-SPRTN-depleted extracts were blotted against SPRTN and two proteasome subunits (PSMA1 and PSMA3). ORC2 was blotted as loading control. (**B**) Extracts from (A) were used to replicate pDPC^2xLead^. DPCs were recovered and monitored like in Figure 1B. (**C**) Extracts from (A) were used to replicate pDPC^2xLead^ and samples were analyzed as in Figure 3B.

**Figure S5.**
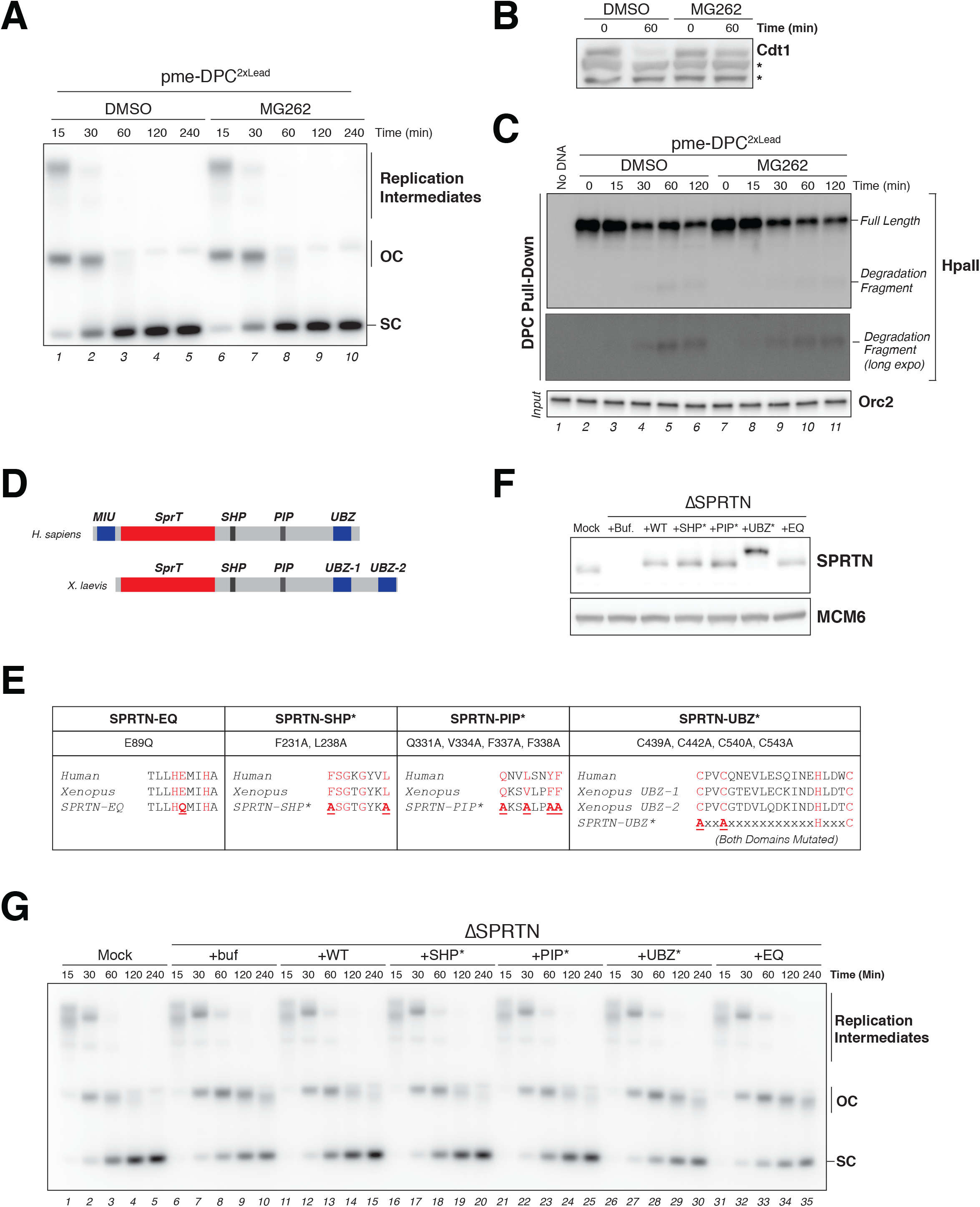
(**A**) pme-DPC^2xLead^ was replicated in egg extracts supplemented with either DMSO (control) or 200 μM of MG262 and analyzed as in Figure 3B. (**B**) Samples from (A) were blotted against CDT1 to control for proteasome inhibition. *denote non-specific bands that are used as loading control. (**C**) Samples from (A) were used to monitor DPC degradation like in Figure 1B using anti-M.HpaII antibody. Two different exposures are shown as indicated. (**D**) Schematic of human and *Xenopus laevis* SPRTN protein. Note that *Xenopus laevis* SPRTN contains a duplication of the C-terminal UBZ domain. (**E**) Alignment of the different functional motifs of human and *Xenopus laevis* SPRTN. Residues mutated to generate EQ, SHP*, PIP* and UBZ* are indicated. (**F**) Mock-depleted and SPRTN-depleted extracts were blotted against SPRTN or MCM6 (loading control). SPRTN-depleted extracts were supplemented with either buffer (+buf), or recombinant FLAG-SPRTN variants. For details about the mutations see (E). (**G**) Extracts from (F) were used to replicate pme-DPC^2xLead^ in the presence of [α-^32^P]dATP. Samples were analyzed by agarose gel electrophoresis as in Figure 3B.

**Figure S6.**
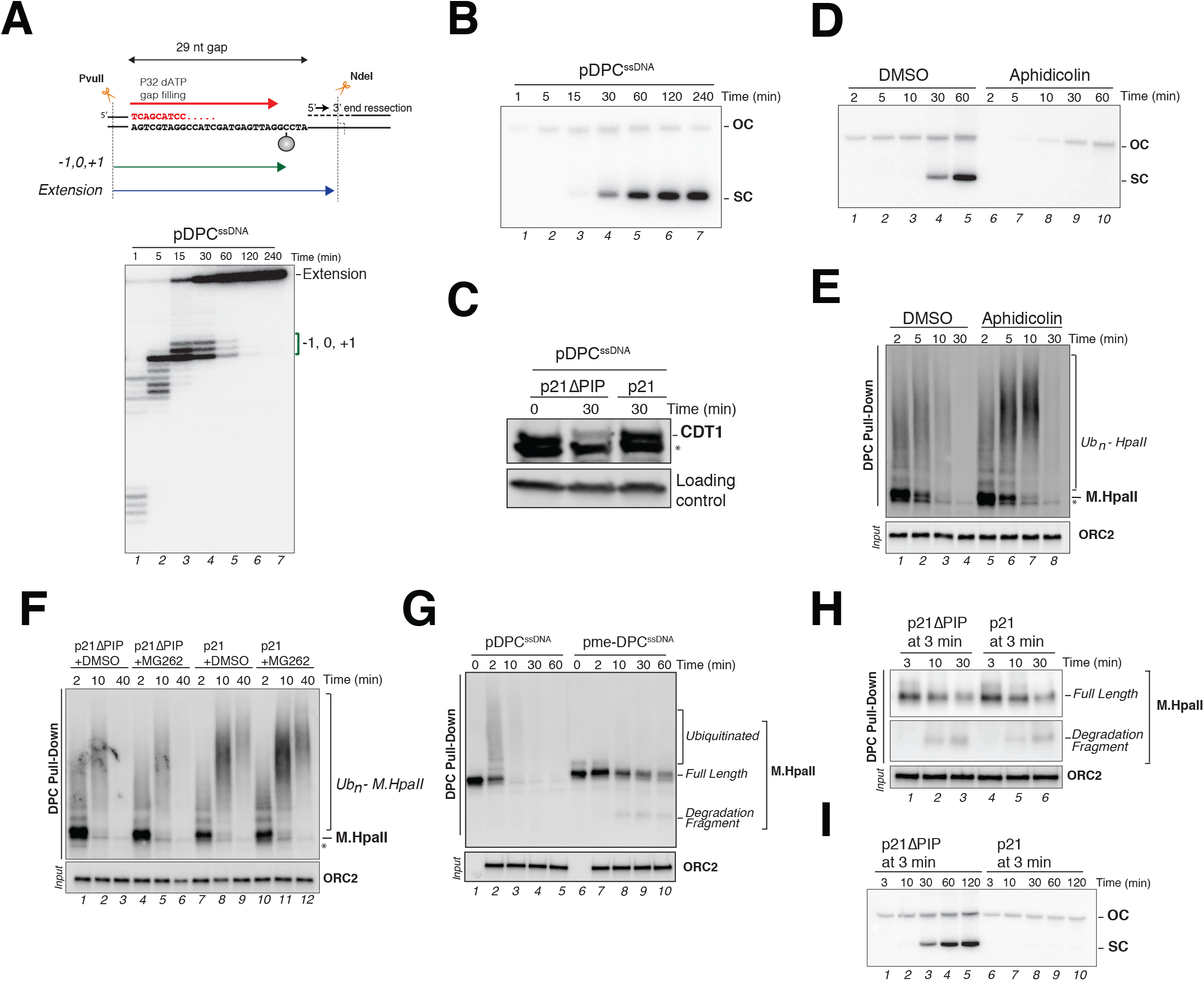
(**A**) Top scheme depicts the 29 nt gap of pDPC^ssDNA^. pDPC^ssDNA^ was incubated in non-licensing egg extracts in the presence of [α-^32^P]dATP. Samples were digested with PvuII and NdeI and separated on a denaturing polyacrylamide gel (bottom autoradiograph). The different extensions products are depicted in the upper scheme. (**B**) Samples from (A) were analyzed by agarose gel electrophoresis as in Figure 3B. (**C**) Extracts from Figure 6C were used to monitor damage-dependent CDT1 degradation. Samples were blotted with a CDT1 or ORC2 (loading control) antibodies. Note that p21 addition, but not p21ΔPIP, blocks CDT1 degradation in accordance with (Arias and Walter, 2006). * denotes a non-specific band. (**D**) pDPC^ssDNA^ was incubated in non-licensing egg extracts supplemented with either DMSO (control) or 200 μM of aphidicolin. Samples were analyzed by agarose gel electrophoresis. (**E**) Samples from (D) were used to monitor DPC ubiquitylation and degradation as in Figure 1B using an M.HpaII antibody. (**F**) pDPC^ssDNA^ was incubated in non-licensing egg extracts supplemented with p21 peptide (p21), p21 peptide with a mutated PIP box (p21ΔPIP) in the presence or absence of 200 μM of MG262. DPCs were recovered and monitored as in Figure 1B. (**G**) pDPC^ssDNA^ or pme-DPC^ssDNA^ were incubated in non-licensing egg extracts. DPCs were recovered and monitored as in Figure 1B. Samples at time 0 were withdrawn prior to incubating plasmids in egg extracts. (**H**) pDPC^ssDNA^ was incubated in non-licensing extracts. At 3 minutes, prior to the detection SPRTN-dependent proteolysis, reactions were supplemented with either p21 or p21ΔPIP peptide. DPCs were monitored as in Figure 1B. (**I**) Samples from (H) were supplemented with [α-^32^P]dATP and analyzed by gel electrophoresis like in Figure 3B. Note that in the presence of the p21 peptide, no SC molecules appear indicative of TLS inhibition.

**Figure S7.**
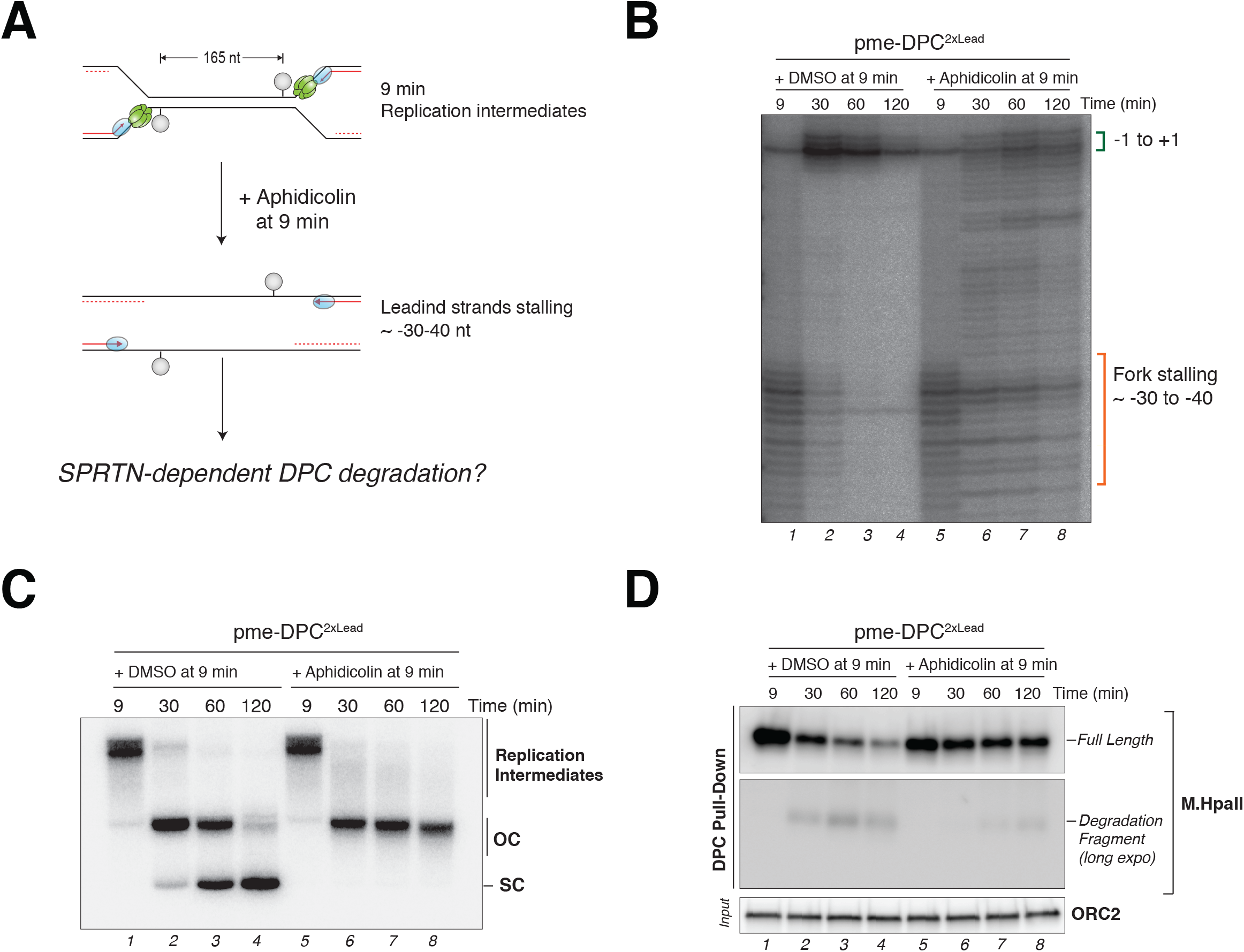
(**A**) Depiction of aphidicolin addition during replication of pme-DPC^2xLead^ at 9 min after the vast majority of forks have reached a DPC located on their leading strand template. (**B**) pme-DPC^2xLead^ was replicated in egg extracts in the presence of [α-^32^P]dATP, and 200 μM of aphidicolin was added at 9 minutes to block polymerase extension as depicted in (A). Samples were analyzed by denaturing polyacrylamide gel electrophoresis following nb.BsmI digestion. Note that nascent leading strand extension from ~-30-40 to the −1, 0, +1 positions is strongly inhibited in the presence of aphidicolin. (**C**) Samples from (B) were analyzed by native agarose gel electrophoresis like in Figure 3B. (**D**) Samples from (B) were used to monitor DPC degradation like in Figure 1B. Note that the residual DPC degradation observed upon aphidicolin treatment likely results from some nascent leading strands escaping inhibition and extending up to the lesion site (B, lanes 6-8).

**Table S1. Protein enrichment on pDPC^2xLead^.** The table summarizes the results from pairwise comparisons using a modified T-test with a permutation-based FDR cut-off implemented in the Perseus frame work (Tyanova et al., 2016). To test for protein enrichment on plasmid substrates a one-sided T-Test was carried out with a FDR<0.01 and S0=4. For each protein in *column A* significant enrichment is indicated in *column E-K* (1: significant, 0: not significant, NA: not detected or not enough valid values). *Column Al-CQ* list the *log2* fold change (columns labelled DIFF) and the associated –LOG_10_(p-value) (columns labelled PVAL). All experiments were measured in quadruplicates. The combined p-value is the product of individual p-values across the time series corrected by random permutation across the entire data matrix (see (Tyanova et al., 2016) for details). **pDPC**, plasmid substrate crosslinked with M.HpaII at both leading strand templates (pDPC^2xLead^) (see Figure 2); **pCTR**, undamaged control plasmid (pCTRL); **Gem**, reactions containing the replication inhibitor geminin; **UB-VS**, reactions containing ubiquitin vinyl sulfone.

**Table S2. Dynamic recruitment of DNA repair factors to pDPC^2xLead^.** The table shows the z-scored log2 LFQ intensities (mean from 4 biochemical replicates) for all quantified proteins *(column A-O).* A subjective chronological order for selected proteins is provided *(column Q*).

